# Interferon lambda signaling to maternal dendritic cells protects against congenital Zika virus infection

**DOI:** 10.64898/2026.01.25.701624

**Authors:** Margaret R. Dedloff, Thomas R. Reinhardt, Tessa-Jonne F. Ropp, Joel Rivera-Cardona, Celeste R. Robles, Rebecca L. Casazza, J. Ashley Ezzell, Michelle S. Itano, Helen M. Lazear

## Abstract

Interferon lambda (IFN-λ, type III IFN) mediates antiviral immunity at anatomic barriers, including the maternal-fetal interface. To investigate the effects of IFN-λ during congenital Zika virus (ZIKV) infection, we infected mice lacking the IFN-⍺β receptor (*Ifnar1*^-/-^) or both the IFN-⍺β and IFN-λ receptors (*Ifnar1*^-/-^*Ifnlr1*^-/-^) at E9 and found that loss of maternal IFN-λ signaling resulted in greater transplacental transmission. We used HiPlex RNAscope on entire gravid uteruses and found that IFN-λ was expressed more proximal to the site of ZIKV infection in *Ifnar1*^-/-^ dams compared to *Ifnar1*^-/-^*Ifnlr1*^-/-^ dams. We performed immunophenotyping of the placenta and uterus by flow cytometry and found a decrease in dendritic cells and NK cells in the uterus of *Ifnar1*^-/-^*Ifnlr1*^-/-^ dams compared to *Ifnar1*^-/-^ dams, but NK cell depletion did not impact fetal infection. Using conditional knockout mice, we identified maternal dendritic cells as the key IFN-λ responsive cell type mediating protection against ZIKV congenital infection.

## INTRODUCTION

Only a subset of pathogens, including Zika virus (ZIKV), rubella virus, and human cytomegalovirus can cross the placental barrier during pregnancy and cause congenital infections^1,2^. Congenital infections can result in a range of birth defects and adverse pregnancy outcomes, such as fetal loss and intrauterine growth restriction (IUGR). Furthermore, the adverse effects of congenital infections can manifest as cognitive and functional disabilities later in childhood even in infants seemingly unaffected at birth^3–5^. ZIKV is a mosquito-borne flavivirus that usually presents with fever and rash, but the 2015-2017 ZIKV outbreak in the Americas revealed the ability of ZIKV to cause congenital Zika syndrome when infection occurs during pregnancy^6–10^. Congenital Zika syndrome is observed in ∼6% of pregnancies with ZIKV infection^18^ and encompasses a spectrum of fetal abnormalities including IUGR, microcephaly, and stillbirth^6–9^.

Immunity at the maternal-fetal interface is a complex balance between protecting the fetus from maternal pathogens and safeguarding against maternal immune-mediated rejection of the semi-allogeneic fetus and placenta^11–13^. The placenta is a fetal-derived tissue that invades the maternal endometrium (the decidua), forming the interface between maternal and fetal tissues during pregnancy. Nutrients are passed from the mother to the fetus through the placental villi, the surface of which is formed by syncytiotrophoblasts, which are specialized cells fused to form a single-cell barrier between maternal and fetal blood. Syncytiotrophoblasts are highly resistant to infection by diverse pathogens, making them an important barrier against fetal infection^14–16^. Conversely, tolerogenic immune mechanisms at the maternal-fetal interface prevent maternal immune rejection of the fetus^17^. This delicate balance between protective and tolerogenic immunity changes over the course of gestation^18^.

Interferon lambda (IFN-λ, type III IFN) is a cytokine that induces a similar antiviral transcriptional program as type I interferons (IFN-αβ) but is especially important at anatomic barriers, including the maternal-fetal interface^19,20^. In contrast to the ubiquitous expression of the IFN-αβ receptor (heterodimer of IFNAR1 and IFNAR2), the IFN-λ receptor (heterodimer of IFNLR1 and IL10Rb) is found predominantly on epithelial cells and a subset of immune cells, including myeloid cells^21–30^. Both IFN-αβ and IFN-λ have immunomodulatory properties, but IFN-λ is thought to be less potent and pro-inflammatory than IFN-αβ^31–39^. During pregnancy, IFN-λ is constitutively secreted by syncytiotrophoblasts^23,40^ and decidual cells^41^. IFN-λ is produced in response to ZIKV infection in trophoblasts in cell culture and primary decidual tissues^40,42^, and treatment of mice with IFN-λ during infection decreases ZIKV viral loads^42,43^. Previously, we demonstrated that IFN-λ acts specifically through signaling in maternal tissues (rather than fetal/placental tissues) following ZIKV infection during pregnancy, but the effects of IFN-λ can be either pathogenic or protective depending on when infection occurs during gestation^44^. IFN-λ reduced ZIKV transplacental transmission when infection occurred later in pregnancy (E9); in contrast, IFN-λ signaling induced fetal pathology when infection occurred earlier in pregnancy (E7). Using poly(I:C) treatment to mimic the IFN-inducing effects of viral infection, we found that the pathogenic effects of IFN-λ during early gestation occurred through IFN-λ signaling in maternal leukocytes^44^, but the maternal cell types responsible for IFN-λ mediated protection against transplacental transmission remain undefined.

Here, we further define the protective role for IFN-λ during congenital ZIKV infection. Using a mouse model of congenital ZIKV infection, we demonstrate that maternal IFN-λ signaling is protective against fetal infection and transplacental transmission. We found that ZIKV infection occurs in fetuses, placentas, and the uterus regardless of maternal IFN-λ signaling, but there is a more robust antiviral response near the site of infection in dams with intact IFN-λ signaling. We found that dams with intact IFN-λ signaling have an increased proportion and number of NK cells at the maternal-fetal interface during congenital ZIKV infection, but these cells were not responsible for IFN-λ mediated protection against fetal infection and transplacental transmission. We show that dendritic cells accumulate at the maternal-fetal interface during congenital ZIKV infection in an IFN-λ dependent manner and we demonstrate that IFN-λ signaling to dendritic cells is required for protection against fetal infection and transplacental transmission. Altogether, our results suggest that IFN-λ plays both immunomodulatory and antiviral roles at the maternal-fetal interface during congenital ZIKV infection and that maternal dendritic cells are key IFN-λ responsive cells at this site.

## RESULTS

### Maternal IFN-λ signaling protects against fetal ZIKV infection in *Ifnar1*^-/-^ mice

Our group and others previously showed that maternal IFN-λ signaling protects against fetal infection when ZIKV infection occurs later in gestation (E9 and E12)^42,44^. However, these experiments used wild-type and *Ifnlr1*^-/-^ mice treated with an IFNAR1-blocking mAb, which is sufficient to promote ZIKV infection, but not virus-induced pathology. To assess the protective effects of IFN-λ in the context of a more robust ZIKV infection, we used pregnancies in which dams and fetuses both lack IFN-αβ signaling (*Ifnar1*^-/-^). We set up crosses such that fetuses (and their associated placentas) all retained IFN-λ signaling either in the context of dams that lacked IFN-λ signaling (*Ifnar1*^-/-^ *Ifnlr1*^-/-^double-knockout) or dams that retained IFN-λ signaling (*Ifnar1*^-/-^), so as to evaluate the effects of IFN-λ signaling specifically in maternal tissues. We evaluated fetal pathology, fetal viral loads, and fetal infection rates in *Ifnar1^-/-^* dams mated with *Ifnar1^-/-^ Ifnlr1^-/-^* sires or *Ifnar1^-/-^ Ifnlr1^-/-^* dams mated with *Ifnar1^-/-^* sires. At 9 days post-mating (E9), pregnant dams were infected with 1000 focus-forming units (FFU) of ZIKV strain FSS13025, and maternal and fetal tissues were harvested 6 days post-infection (dpi) (E15) to measure viral loads and assess fetal pathology (Figure 1A). We found no difference in maternal spleen viral loads between *Ifnar1^-/-^* dams and *Ifnar1^-/-^ Ifnlr1^-/-^* dams (Figure 1B), but fetuses from dams lacking IFN-λ signaling exhibited significantly higher viral loads compared to dams with intact IFN-λ signaling (median fetal viral loads 3.41 vs. 5.06 Log_10_ copies/mL, *P* < 0.0001, Figure 1C). Additionally, we found that dams lacking IFN-λ signaling had increased rates of fetal infection compared to dams with intact IFN-λ signaling (97% vs. 87% of fetuses positive for ZIKV RNA, *P* < 0.05, Figure 1D). We also measured infectious viral titers in the fetuses by plaque assay and found that dams lacking IFN-λ signaling had significantly higher fetal viral loads (median 3.32 vs. 3.41 Log_10_ plaque forming units/g, *P* < 0.05, Figure S1A) and fetal infection rates (36% vs. 57% of fetuses positive for virus, *P* < 0.05, Figure S1B) compared to dams with intact IFN-λ signaling, consistent with RT-qPCR results. All fetuses were weighed and photographed to evaluate two measurements of fetal pathology: intrauterine growth restriction, defined as fetal weights below one standard deviation of the mean of fetuses from mock-infected mice, and resorptions, which were defined as implantation sites with no discernable fetal or placental structures (Figure 1E-G). We found that there was a significant decrease in fetal weights from dams with intact IFN-λ signaling, but no differences in fetal pathology between dams with intact IFN-λ signaling and dams that lack IFN-λ signaling. Overall resorption and intrauterine growth restriction rates were low, likely due to the lack of fetal IFN-αβ signaling in this experimental design^44,45^.

**Figure 1:**
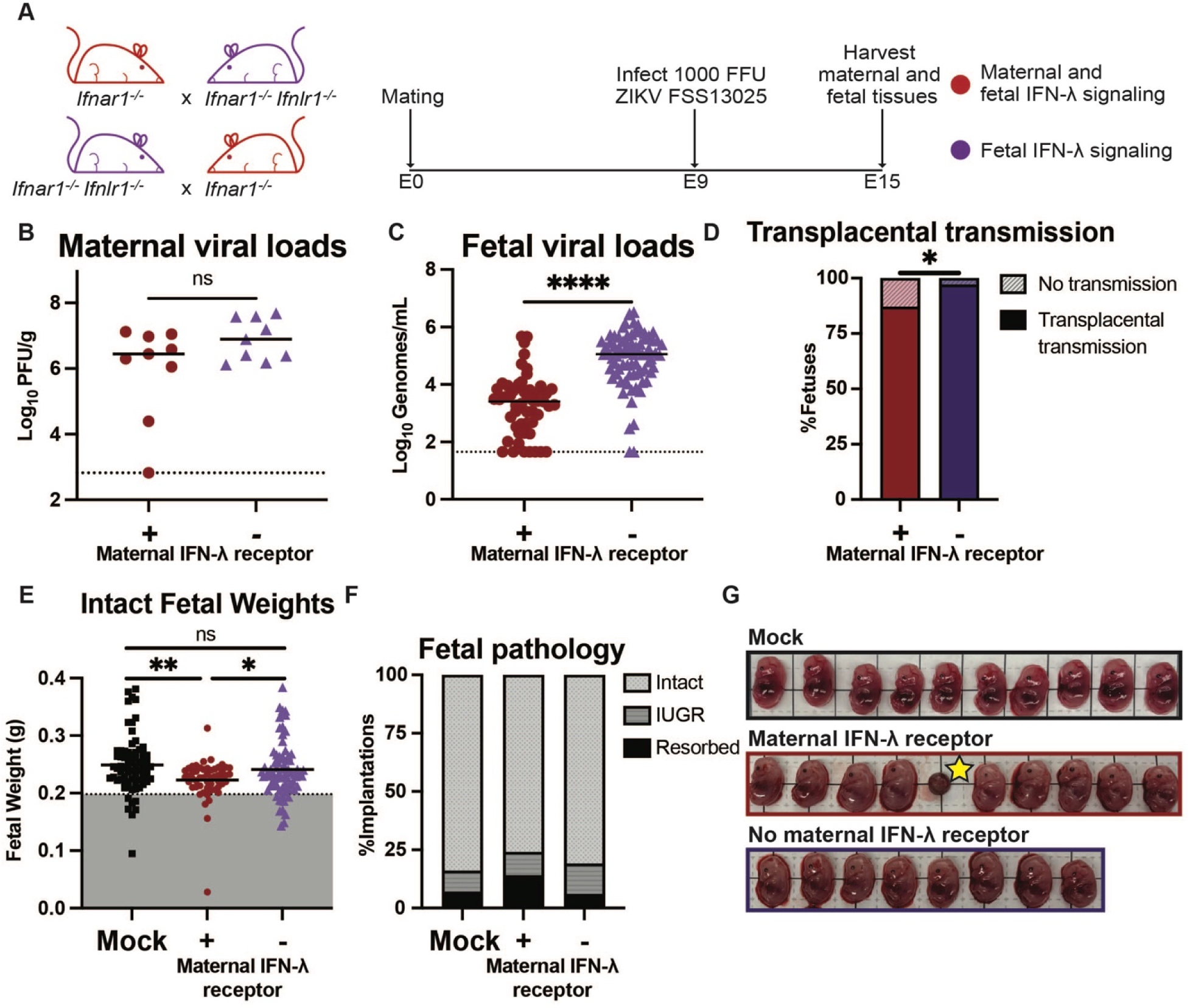
Maternal IFN-λ signaling protects against fetal ZIKV infection in *Ifnar1*^-/-^ mice. (A) Mating and infection timeline. Dams from *Ifnar1^-/-^* x *Ifnar1^-/-^ Ifnlr1^-/-^* or *Ifnar1^-/-^ Ifnlr1^-/-^* x *Ifnar1^-/-^* crosses were infected at day 9 post-mating (E9) with ZIKV FSS13025 by subcutaneous injection in the footpad. Maternal and fetal tissues were harvested at E15. (B) Viral loads in the maternal spleen were measured by plaque assay. Each data point represents one dam. (C) Fetal viral loads were measured by RT-qPCR. Each data point represents one fetus. (D) Transplacental transmission was calculated as percent of fetuses with viral loads above the limit of detection. (E) Intact fetuses were weighed. Dotted line indicates 1 standard deviation below the mean of fetuses from mock-infected dams; fetal weights below this threshold were classified as having intrauterine growth restriction (IUGR). Fetal weights were compared by *t*-test (**, *P* < 0.01; *, *P* <0.05; ns, no significant difference). (F) Proportions of implantation sites exhibiting IUGR or resorption. (G) Representative images of one pregnancy from each cross. The star indicates resorption. Each square in the grid is 1cm. Results are combined from 62 fetuses from 9 *Ifnar1*^-/-^ dams, 82 fetuses from 10 *Ifnar1^-/-^ Ifnlr1^-/-^* dams, and 78 fetuses from 9 uninfected dams from 11 independent experiments. Viral loads were compared by Mann-Whitney test, proportions by Cochran Armitage test (***, *P* < 0.0001; *, *P* < 0.05; ns, no significant difference).

### IFN-λ protects against fetal infection regardless of infection duration

Previously, we found that maternal IFN-λ signaling can have either protective or pathogenic effects depending on when infection occurred during gestation^44^. Notably, these results were based on fetuses harvested at E15, meaning that when infections occurred at E7, fetuses were harvested 8 dpi, compared to 6 dpi for infections at E9 (Figure S2A), leaving the possibility that the pathogenic effects of IFN-λ with E7 infection only become evident as the infection progressed. To address this question, we evaluated viral loads, infection rates, and pathology of fetuses harvested from dams at E15 (6 dpi) or E17 (8 dpi). We mated dams with intact IFN-λ signaling (*Ifnar1*^-/-^) with sires lacking IFN-λ signaling (*Ifnar1^-/-^ Ifnlr1^-/-^*) or the reverse mating scheme. Dams were infected at E9 with 1000 FFU of ZIKV. We found that dams lacking IFN-λ signaling had significantly higher fetal viral loads compared to dams with intact IFN-λ signaling at both 6 dpi (median 3.47 vs. 6.80 Log_10_ copies/mL, *P* < 0.0001) and 8 dpi (median 4.14 vs. 5.49 Log_10_ copies/mL, *P* < 0.0001, Figure S2B). Furthermore, at both timepoints, dams lacking IFN-λ signaling had significantly higher rates of transplacental transmission compared to dams with intact IFN-λ signaling (6 dpi 84% vs. 100%, *P* < 0.01; 8 dpi 84% vs. 98%, *P* < 0.05, Figure S2C). Maternal IFN-λ signaling did not impact resorption rate (Figure S2D). Altogether, these results show that maternal IFN-λ signaling protects against transplacental transmission after E9 ZIKV infection, regardless of infection duration.

### ZIKV infection occurs in fetuses, placentas, and the uterus

Because the architecture of the maternal-fetal interface is complex and unique, determining where viral infection occurs is critical to understanding the pathogenic mechanisms of congenital infections^1,46,47^. To determine where ZIKV infection is occurring at the maternal-fetal interface, we mated *Ifnar1^-/-^* dams with *Ifnar1^-/-^Ifnlr1^-/-^* sires and *Ifnar1^-/-^ Ifnlr1^-/-^* dams with *Ifnar1^-/-^* sires, infected dams with ZIKV at E9, and harvested tissues at E13 to evaluate the spatial localization of ZIKV RNA in the gravid uterus using HiPlex RNAscope. In separate pregnancies, we measured viral loads by RT-qPCR (Figure 2A). We detected ZIKV in fetuses, placentas, and the uterus from both *Ifnar1^-/-^* and *Ifnar1^-/-^ Ifnlr1^-/-^* dams (Figure 2B, Figure S3). Dams lacking IFN-λ signaling exhibited significantly higher viral loads in fetuses (median 7.11 vs. 6.18 Log_10_ copies/g, *P* < 0.0001) and placentas (median 8.20 vs. 7.87 Log_10_ copies/g, *P* < 0.001) compared to dams with intact IFN-λ signaling (Figure 2C-D), but there was no significant difference in viral loads in the uterus (Figure 2E). These findings demonstrate that maternal IFN-λ signaling does not impact infection in maternal tissues but protects against ZIKV transmission to the placenta and fetus.

**Figure 2:**
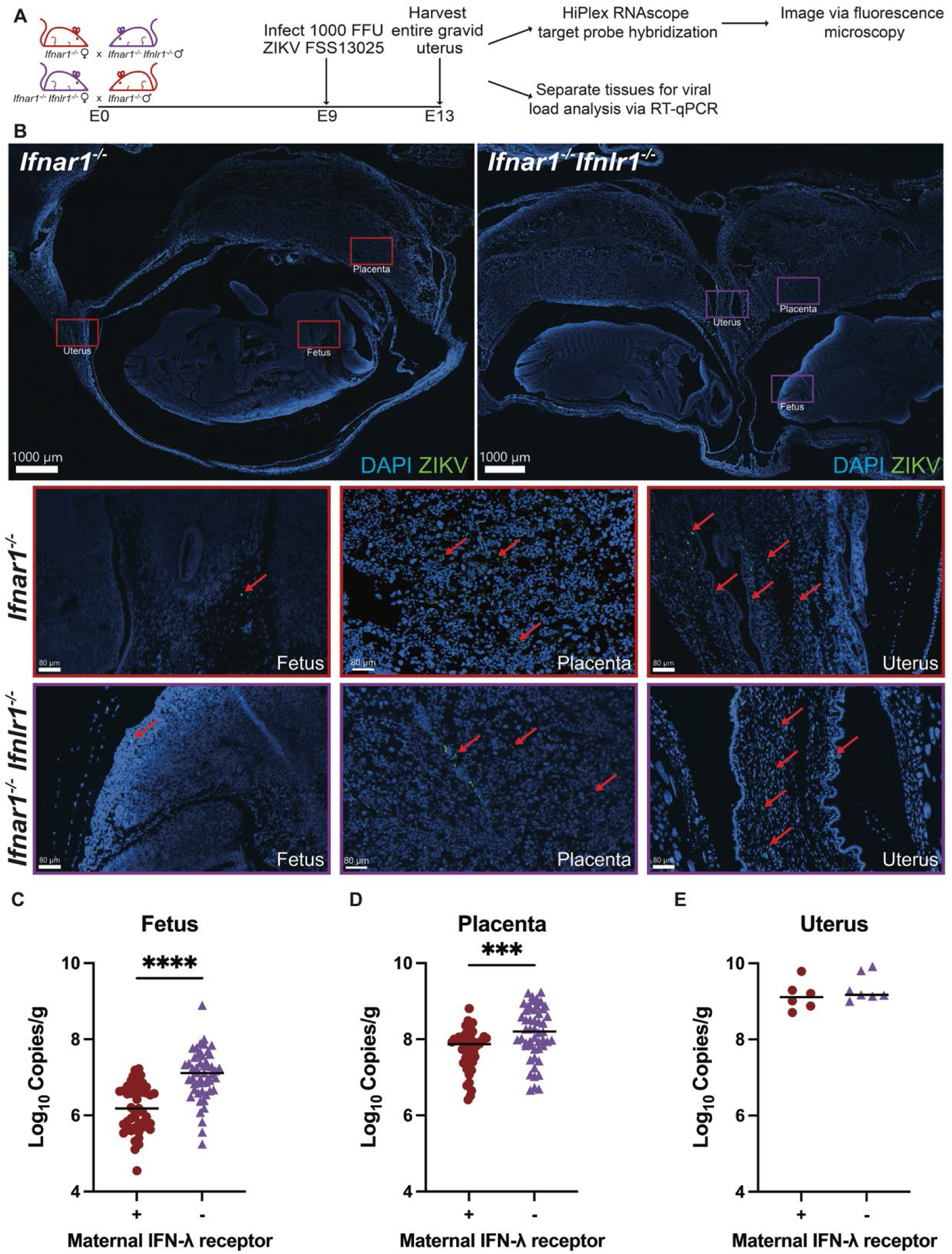
ZIKV infection occurs in fetuses, placentas and the uterus. (A) Mating and infection timeline. Dams from *Ifnar1^-/-^* x *Ifnar1^-/-^ Ifnlr1^-/-^* or *Ifnar1^-/-^ Ifnlr1^-/-^* x *Ifnar1^-/-^* crosses were infected at E9 with ZIKV FSS13025 by subcutaneous injection in the footpad. Maternal and fetal tissues were harvested at E13. (B) Representative images of portions of implantation sites. Sections of entire gravid uteruses (2 *Ifnar1*^-/-^ dams and 2 *Ifnar1^-/-^ Ifnlr1^-/-^* dams per group, collected in independent experiments) were probed for ZIKV RNA (green) and stained with DAPI (blue). Arrows indicate ZIKV staining. Scale bars in top row of images are 1000µm and in the bottom rows of images are 80µm. (C to E) Viral loads in fetuses, placentas, and uterus were measured by RT-qPCR. Each data point represents one fetus (C), one placenta (D), or one dam (E). Results are combined from 52 fetuses from 6 *Ifnar1*^-/-^ dams and 58 fetuses from 7 *Ifnar1^-/-^ Ifnlr1^-/-^* dams from 7 independent experiments. Viral loads were compared by Mann-Whitney test (****, *P* < 0.0001; ***, *P* < 0.001).

### Maternal IFN-λ signaling increases IFN-λ expression at the site of infection

To investigate the spatial relationship between ZIKV infection and the IFN-λ response, we mated *Ifnar1^-/-^* dams with *Ifnar1^-/-^ Ifnlr1^-/-^* sires and *Ifnar1^-/-^ Ifnlr1^-/-^* dams with *Ifnar1^-/-^* sires, infected dams with ZIKV at E9, harvested the entire gravid uterus at E13 and probed for ZIKV and *Ifnl2/3* RNA by HiPlex RNAscope. Using Imaris Microscopy Image Analysis Software, we assigned spots for ZIKV and *Ifnl2/3* and color-coded *Ifnl2/3* spots based on the distance to the nearest ZIKV spot (Figure 3A). We found that *Ifnl2/3* spots were closer to ZIKV spots in dams with intact IFN-λ signaling compared to dams lacking IFN-λ signaling (Figure 3B). To determine if the proximity of *Ifnl2/3* expression to ZIKV infection reflected changes in antiviral gene expression across the entire placenta, we mated *Ifnar1^-/-^* dams with *Ifnar1^-/-^ Ifnlr1^-/-^* sires and *Ifnar1^-/-^ Ifnlr1^-/-^* dams with *Ifnar1^-/-^*sires, infected dams with ZIKV at E9, harvested placentas at E15, and detected *Ifnl3, Ifnb1,* and *Ifit1* expression by RT-qPCR (Figure 3C-E). Gene expression was compared to *Actb* and normalized to the baseline gene expression in placentas from uninfected dams. We found no effect of maternal IFN-λ signaling on induction of *Ifnl3, Ifnb1,* or *Ifit1* in placenta homogenates, highlighting the importance of the spatial relationship between the IFN-λ mediated antiviral response and control of ZIKV infection at the maternal-fetal interface.

**Figure 3:**
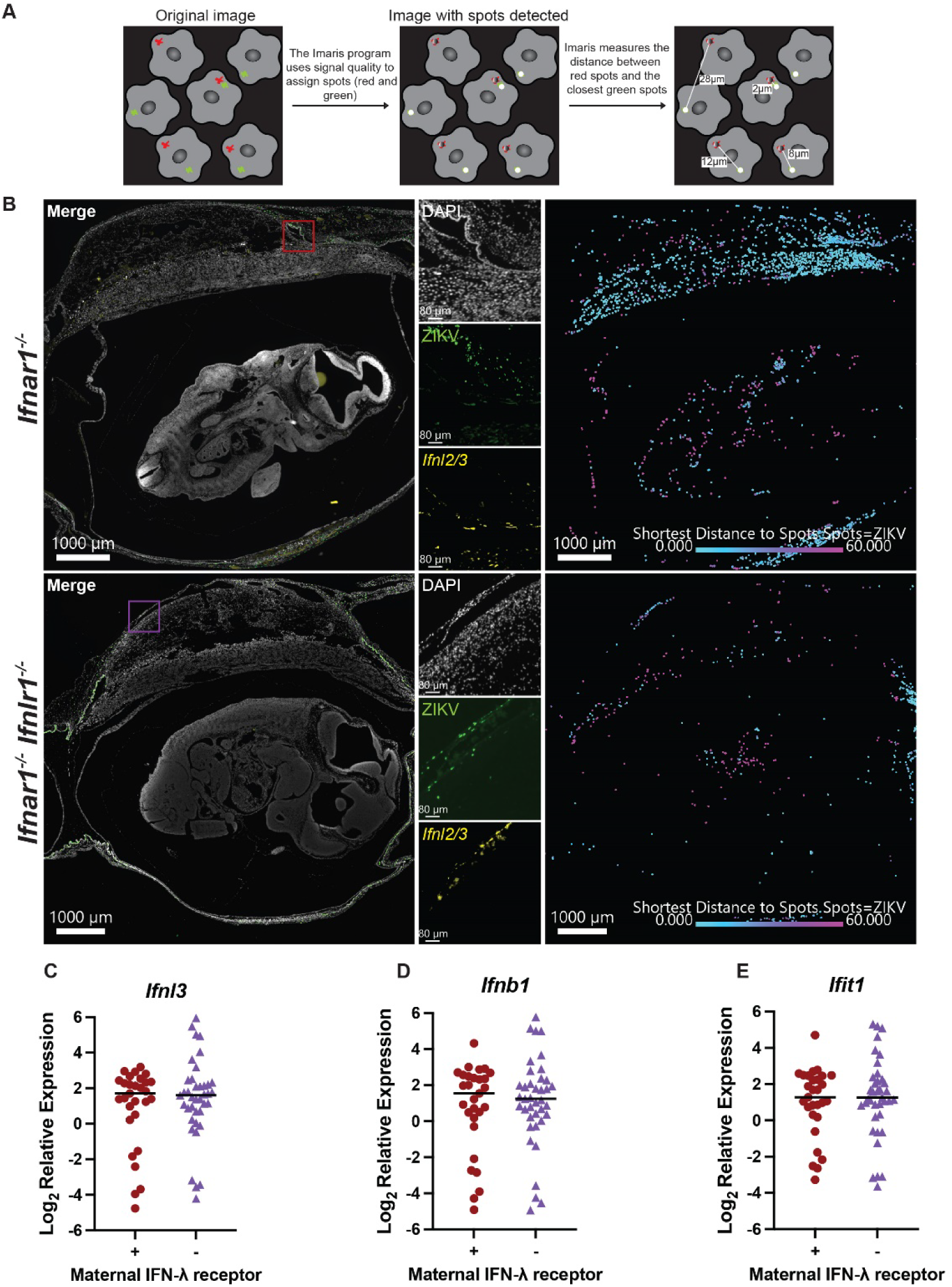
Maternal IFN-λ signaling increases IFN-λ expression at the site of infection. (A) Image analysis pipeline. Spots are assigned to signal in Imaris based on quality. Imaris measures the distance between spots. (B) Representative images of implantation sites. Dams from *Ifnar1^-/-^* x *Ifnar1^-/-^ Ifnlr1^-/-^* or *Ifnar1^-/-^ Ifnlr1^-/-^* x *Ifnar1^-/-^* crosses were infected at E9 with ZIKV FSS13025 by subcutaneous injection in the footpad. Gravid uteruses were harvested at E13. Sections of entire gravid uteruses (2 *Ifnar1*^-/-^ dams and 2 *Ifnar1^-/-^ Ifnlr1^-/-^* dams per group, collected in independent experiments) were probed for ZIKV RNA (green) and *Ifnl2/3* RNA (yellow) and stained with DAPI (white). Spots were assigned to ZIKV and *Ifnl2/3* signal. IFN-λ spots are colored based on their distance to the closest ZIKV spot (right most images of B). Scale bars in zoomed images are 80µm. (C to E) Dams from *Ifnar1^-/-^* x *Ifnar1^-/-^ Ifnlr1^-/-^* or *Ifnar1^-/-^ Ifnlr1^-/-^* x *Ifnar1^-/-^* crosses were infected at E9 with ZIKV FSS13025 by subcutaneous injection in the footpad. Maternal and fetal tissues were harvested at E15. *Ifnl3* (C), *Ifnb1* (D), and *Ifit1* (E) expression was measured in placentas by RT-qPCR. Gene expression was compared to *Actb* and normalized to the baseline gene expression in placentas from uninfected dams. Each data point represents one placenta. Results are combined from 21 placentas from 3 uninfected *Ifnar1^-/-^* dams, 18 placentas from 2 uninfected *Ifnar1^-/-^ Ifnlr1^-/-^* dams, 29 placentas from 5 ZIKV infected *Ifnar1^-/-^* dams, and 39 placentas from 5 ZIKV infected *Ifnar1^-/-^ Ifnlr1^-/-^* dams collected in 8 independent experiments.

### Maternal IFN-λ signaling induces NK cell accumulation in the uterus during congenital ZIKV infection

To determine how maternal IFN-λ signaling regulates immune cell populations at the maternal-fetal interface during congenital ZIKV infection, we used flow cytometry and a broad immunophenotyping panel (Figure S4-S6). We mated *Ifnar1^-/-^* dams with *Ifnar1^-/-^ Ifnlr1^-/-^* sires and *Ifnar1^-/-^ Ifnlr1^-/-^* dams with *Ifnar1^-/-^* sires, infected dams with ZIKV at E9, and harvested uteruses and placentas at E15. Comparing *Ifnar1^-/-^* dams to *Ifnar1^-/-^ Ifnlr1^-/-^* dams, we observed some differences in leukocyte populations in placentas, but these differences were either in number of cells or percentage, but not both, potentially due to the large variation among placentas in live cell yield and total leukocyte numbers (Figure S4). In the uterus, we found no significant difference in T cells (CD4+, CD8+), B cells, macrophages, or monocytes between *Ifnar1^-/-^* dams and *Ifnar1^-/-^Ifnlr1^-/-^* dams; we found a significant decrease in number (but not percentage) of ɣδ T cells (Figure S5). However, we found that the uteruses of dams lacking IFN-λ signaling had a significantly lower percentage (mean 10 % vs. 15%, *P* < 0.01) and number (mean 876 vs. 3039, *P* < 0.001, Figure 4A-C) of (natural killer) NK cells compared to dams with intact IFN-λ signaling. In contrast, dams lacking IFN-λ signaling exhibited a lower percentage, but not number of NK cells in placentas (mean 2.8% vs. 1.7%, *P* < 0.0001; mean 94 cells vs. 90 cells, Figure 4D-F). Altogether, these results suggest that maternal IFN-λ signaling promotes NK cell accumulation in the uterus during congenital ZIKV infection.

**Figure 4:**
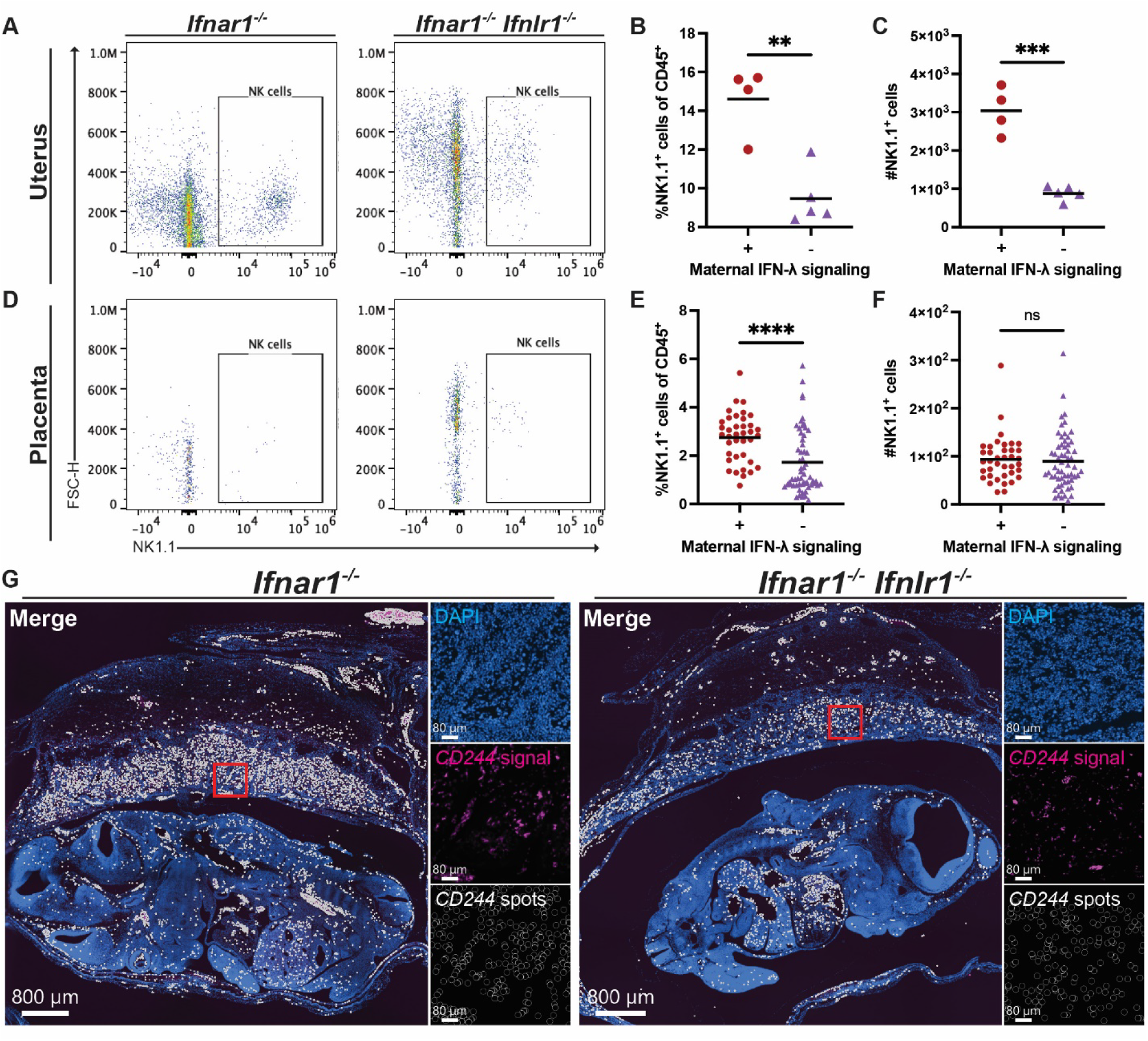
Maternal IFN-λ signaling induces NK cell accumulation in the uterus during congenital ZIKV infection. Dams from *Ifnar1^-/-^* x *Ifnar1^-/-^ Ifnlr1^-/-^* or *Ifnar1^-/-^ Ifnlr1^-/-^* x *Ifnar1^-/-^* crosses were infected at E9 with ZIKV FSS13025 by subcutaneous injection in the footpad. (A) Gravid uteruses were harvested at E15 and analyzed by flow cytometry (4 *Ifnar1*^-/-^ dams and 5 *Ifnar1^-/-^ Ifnlr1^-/-^*dams collected in 4 independent experiments). Representative flow cytometry plots of leukocytes stained for NK1.1. (B-C) Frequency and number of NK cells (NK1.1^+^ out of CD45^+^) in uteruses from *Ifnar1*^-/-^ and *Ifnar1^-/-^ Ifnlr1^-/-^* dams. Groups were compared by unpaired *t*-test (***, *P* < 0.001; **, *P* < 0.01). (D) Placentas were harvested at E15 and analyzed by flow cytometry (38 placentas from 5 *Ifnar1*^-/-^ dams and 59 placentas from 8 *Ifnar1^-/-^ Ifnlr1^-/-^* dams collected in 6 independent experiments). Representative flow cytometry plots of leukocytes stained for NK1.1. (E-F) Frequency and number of NK cells (NK1.1^+^ out of CD45^+^) in placentas from *Ifnar1*^-/-^ and *Ifnar1^-/-^Ifnlr1^-/-^* dams. Groups were compared by unpaired *t*-test (****, *P* < 0.0001; ns, no significant difference). (G) Representative images of implantation sites. Gravid uteruses were harvested at E13. Sections of entire gravid uteruses (2 *Ifnar1*^-/-^ dams and 2 *Ifnar1^-/-^ Ifnlr1^-/-^* dams per group, collected in independent experiments) were probed for *CD244* RNA (magenta), indicating NK cells, and stained with DAPI (blue). Spots were assigned based on the quality of the *CD244* signal (white). Scale bars in zoomed images are 80µm.

**Figure 5:**
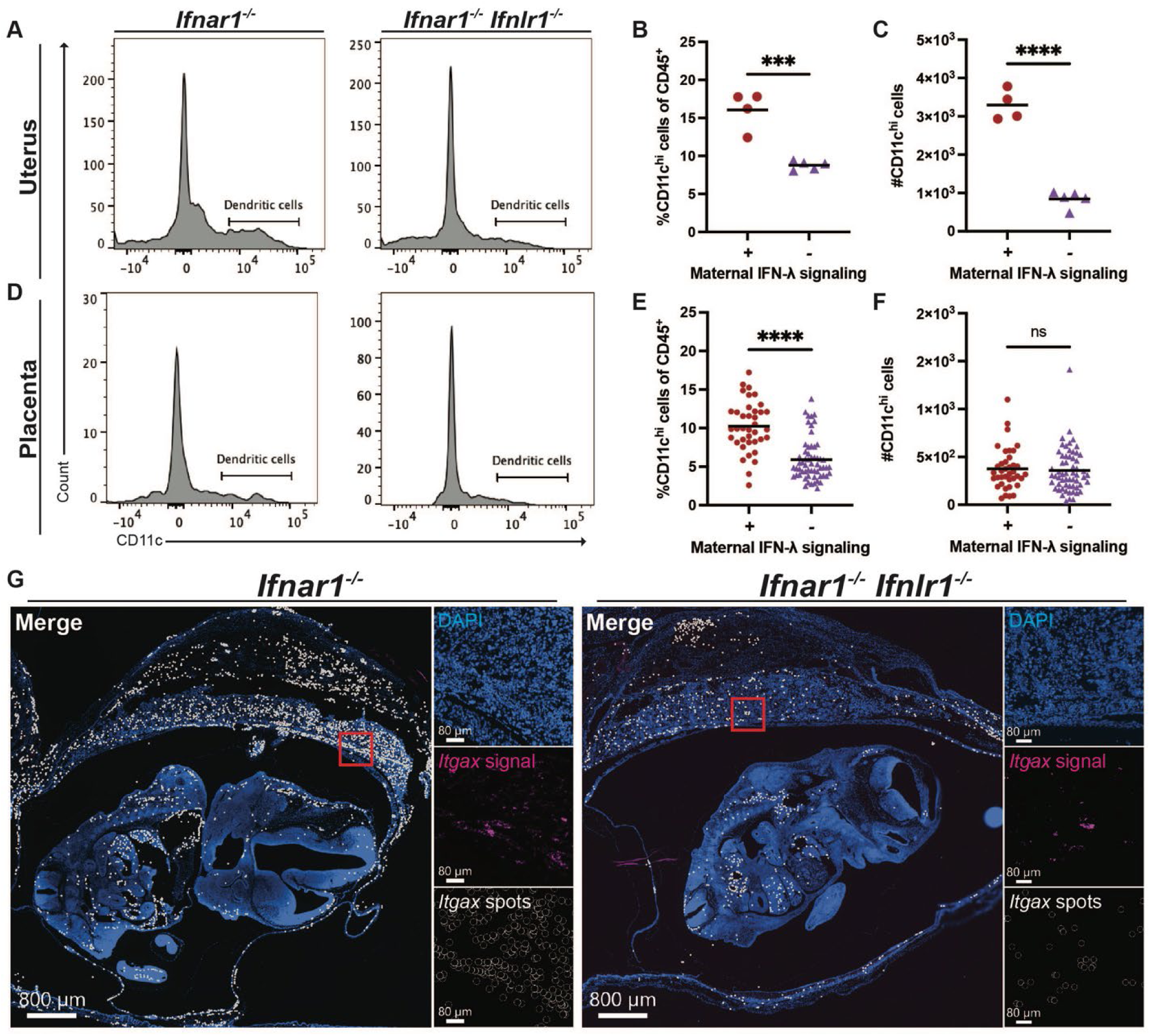
IFN-λ signaling promotes dendritic cell infiltration to the uterus during congenital ZIKV infection. Dams from *Ifnar1^-/-^* x *Ifnar1^-/-^ Ifnlr1^-/-^* or *Ifnar1^-/-^ Ifnlr1^-/-^* x *Ifnar1^-/-^* crosses were infected at E9 with ZIKV FSS13025 by subcutaneous injection in the footpad. (A) Gravid uteruses were harvested at E15 and analyzed by flow cytometry (4 *Ifnar1*^-/-^ dams and 5 *Ifnar1^-/-^ Ifnlr1^-/-^* dams collected in 4 independent experiments). Representative flow cytometry plots of leukocytes stained for CD11c in the uterus. (B-C) Frequency and number of dendritic cells (CD11c^hi^ out of CD45^+^) in uteruses from *Ifnar1*^-/-^ and *Ifnar1^-/-^ Ifnlr1^-/-^* dams. Groups were compared by unpaired *t*-test (****, *P* < 0.0001; ***, *P* < 0.001). (D) Placentas were harvested at E15 and analyzed by flow cytometry (38 placentas from 5 *Ifnar1*^-/-^ dams and 59 placentas from 8 *Ifnar1^-/-^ Ifnlr1^-/-^* dams collected in 6 independent experiments). Representative flow cytometry plots of leukocytes stained for CD11c in placentas. (E-F) Frequency and number of dendritic cells (CD11c^hi^ out of CD45^+^) in placentas from *Ifnar1*^-/-^ and *Ifnar1^-/-^ Ifnlr1^-/-^* dams. Groups were compared by unpaired *t*-test (****, *P* < 0.0001; ns, no significant difference). (G) Representative images of implantation sites. Gravid uteruses were harvested at E13. Sections of entire gravid uteruses (2 *Ifnar1*^-/-^ dams and 2 *Ifnar1^-/-^ Ifnlr1^-/-^* dams per group, collected in independent experiments) were probed for *Itgax* RNA (pink), indicating dendritic cells, and stained with DAPI (blue). Spots were assigned based on the quality of the *Itgax* signal (white). Scale bars in zoomed images are 80µm.

To determine if IFN-λ dependent NK cell accumulation at the maternal-fetal interface contributes to protection against fetal infection, we mated *Ifnar1^-/-^* dams with *Ifnar1^-/-^ Ifnlr1^-/-^* sires, infected with ZIKV at E9 and depleted NK cells. We treated dams intraperitoneally with 100µg of ⍺NK1.1 antibody at E9, E11, and E13, and harvested maternal and fetal tissues at E15 to measure fetal viral loads. We evaluated the efficiency of NK cell depletion systemically and at the maternal-fetal interface using flow cytometry (Figure S7A). NK cells were efficiently depleted from both whole blood (mean 3.0% vs. 22.6% of leukocytes, *P* < 0.01, Figure S7B) and the spleen (mean 0.1% vs. 1.7% of leukocytes, *P* < 0.01, Figure S7C). NK cells were also efficiently depleted in the uterus (mean 3.3% vs. 13.4% of leukocytes, ns) and placentas (mean 0.2% vs. 4.1% of leukocytes, *P* < 0.0001) in dams that received the ⍺NK1.1 antibody (Figure S7D-E).

Despite efficient depletion of NK cells systemically and in tissues at the maternal-fetal interface, we did not observe any difference in fetal viral loads (median 5.40 vs. 5.71 Log_10_ copies/mL, Figure 7G), suggesting that NK cells are not required for IFN-λ mediated protection against ZIKV fetal infection. To determine whether maternal IFN-λ signaling affects NK cell localization at the maternal-fetal interface, we used RNAscope to compare *CD244* spots (indicating NK cells) in *Ifnar1*^-/-^ vs. *Ifnar1*^-/-^ *Ifnlr1*^-/-^ dams infected with ZIKV at E9 and harvested at E13 (Figure 4G). We found that *CD244* signal was mainly found in the uterus and placentas, with less signal in the fetuses and this localization did not depend on maternal IFN-λ signaling. However, the total amount of *CD244* signal was reduced in the absence of IFN-λ signaling, consistent with the reduction in NK cells observed by flow cytometry.

### IFN-λ signaling promotes dendritic cell infiltration to the uterus during congenital ZIKV infection

When assessing immune cell populations at the maternal-fetal interface, we found that the uteruses of dams lacking IFN-λ signaling had fewer dendritic cells (CD45^+^, CD11c^hi^) compared to dams with intact IFN-λ signaling as both percentage of leukocytes (mean 16.1% vs. 8.8%, *P* < 0.001) and number (mean 3292 vs. 844 cells, *P* < 0.0001, Figure 5A-C). In the placentas we found an IFN-λ dependent increase in dendritic cells as percentage of leukocytes (10.2% vs. 5.9%, *P* < 0.0001), but no significant difference in dendritic cell number (mean 375 vs. 359 cells, Figure 5D-F). We used RNAscope to determine the spatial distribution of dendritic cells at the maternal-fetal interface. We harvested tissues at E13, probed for *Itgax* (CD11c), and assigned *Itgax+* spots in Imaris (Figure 5G). As with *CD244*, we observed *Itgax* signal largely in the uterus and placentas, with some signal detected in the fetuses. Furthermore, while the spatial distribution of *Itgax* signal did not change between *Ifnar1^-/-^* and *Ifnar1^-/-^ Ifnlr1^-/-^* dams, we found a decrease in *Itgax* signal in *Ifnar1^-/-^ Ifnlr1^-/-^* dams compared to *Ifnar1^-/-^* dams. Altogether, these results suggest that maternal IFN-λ signaling promotes dendritic cell accumulation at the maternal-fetal interface during congenital ZIKV infection.

### Maternal IFN-λ signaling to dendritic cells protects against transplacental transmission

To determine the IFN-λ-responsive maternal cell type that mediates protection against transplacental transmission, we used conditional knockout dams that lack IFN-λ signaling ubiquitously (*Actin*-Cre-*Ifnlr1^-/-^*), in myeloid cells (*LysM*-Cre-*Ifnlr1^-/-^*), in neutrophils (*Mrp8*-Cre-*Ifnlr1^-/-^*), or dendritic cells (*CD11c*-Cre-*Ifnlr1^-/-^*). These mice were bred as Cre hemizygotes, which results in litters that are 50% Cre+ and 50% Cre-. Experiments were conducted blinded as dams were genotyped following harvesting to allow retrospective comparison of Cre+ and Cre-dams. Conditional knockout dams were mated with wild-type sires and infected with ZIKV at E9, one day after administration of 2mg of IFNAR1-blocking mAb MAR1-5A3. Maternal and fetal tissues were harvested at E15 (Figure 6A). We found no difference in maternal spleen viral loads between Cre-dams and any *Ifnlr1* conditional knockout dams (Figure 6B), consistent with our findings that IFN-λ signaling does not impact ZIKV viral loads in maternal tissues (Figure 1B). *Actin*-Cre-*Ifnlr1^-/-^*dams had significantly higher fetal viral loads (median 4.07 vs. 1.94 Log_10_ copies/mL, *P* < 0.0001, Figure 6C) and fetal infection rates (86% vs. 56%, *P* < 0.0001, Figure 6D) compared to Cre-dams, consistent with what we observe in mice on an *Ifnar1^-/-^* background (Figure 1). Altogether, these results further support a role for maternal IFN-λ signaling in controlling fetal infection.

**Figure 6:**
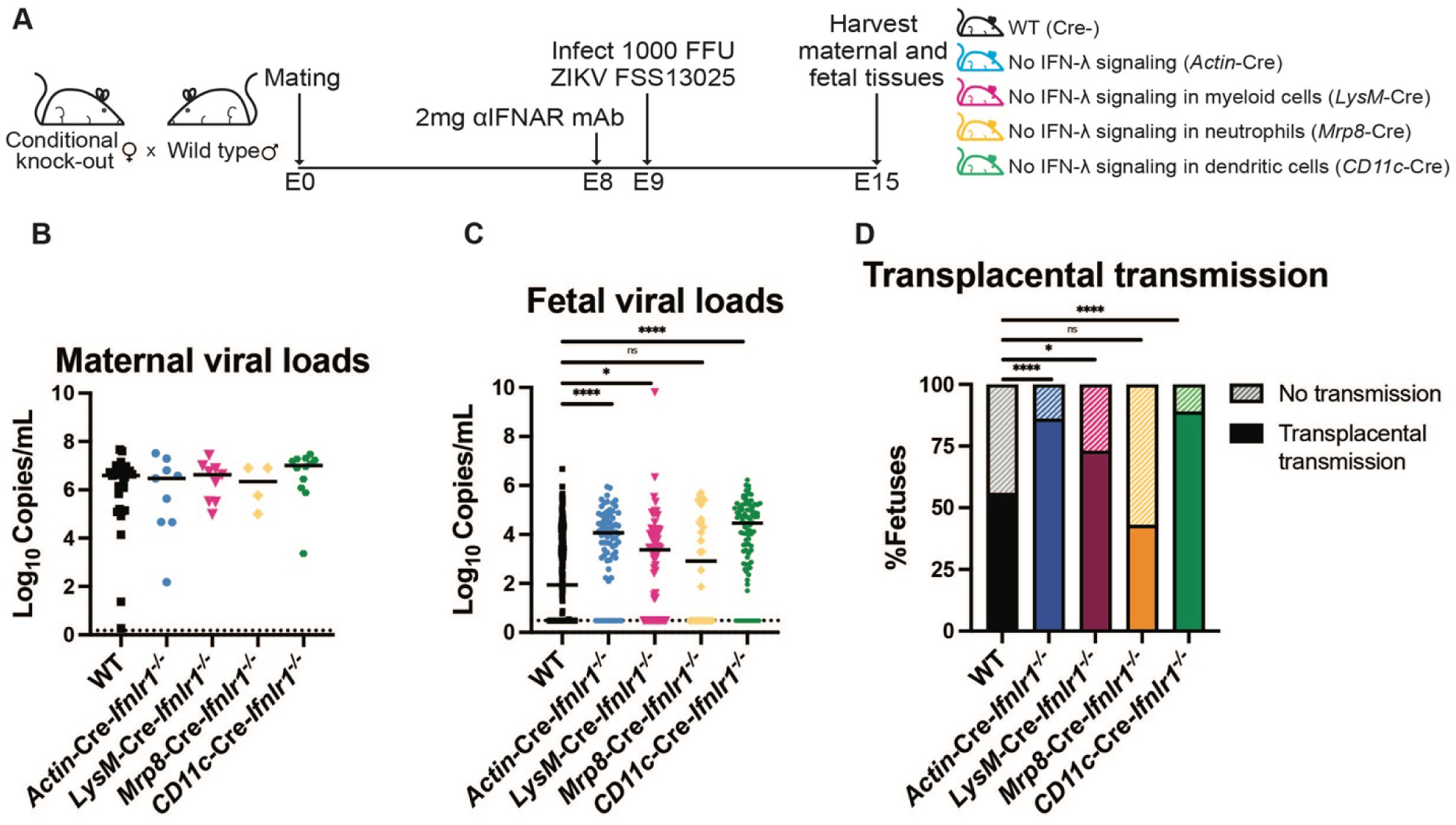
Maternal IFN-λ signaling to dendritic cells protects against transplacental transmission. (A) Mating and infection timeline. Dams from conditional knockout x wild-type crosses (265 fetuses from 34 WT (Cre-) dams, 82 fetuses from 10 *Actin-*Cre-*Ifnlr1^-/-^* dams, 96 fetuses from 12 *LysM-*Cre-*Ifnlr1^-/-^* dams, 30 fetuses from 4 *Mrp8-*Cre-*Ifnlr1^-/-^* dams, and 86 fetuses from 11 *CD11c-*Cre-*Ifnlr1^-/-^* dams) were treated with 2mg of anti-IFNAR1 mAb at E8 and infected at E9 with ZIKV FSS13025 by subcutaneous injection in the footpad. Maternal and fetal tissues were harvested at E15. (B and C) Viral loads in maternal spleens and fetuses were measured by RT-qPCR. Each data point represents one dam (B) or one fetus (C). Cre+ groups were compared to Cre-by Mann-Whitney test. (****, *P* < 0.0001; *, *P* < 0.05; ns, no significant difference). (D) Transplacental transmission was calculated as percent of fetuses with viral loads above the limit of detection. Groups were compared by Cochran Armitage test (****, *P* < 0.0001; *, *P* < 0.05; ns, no significant difference).

**Figure 7.**
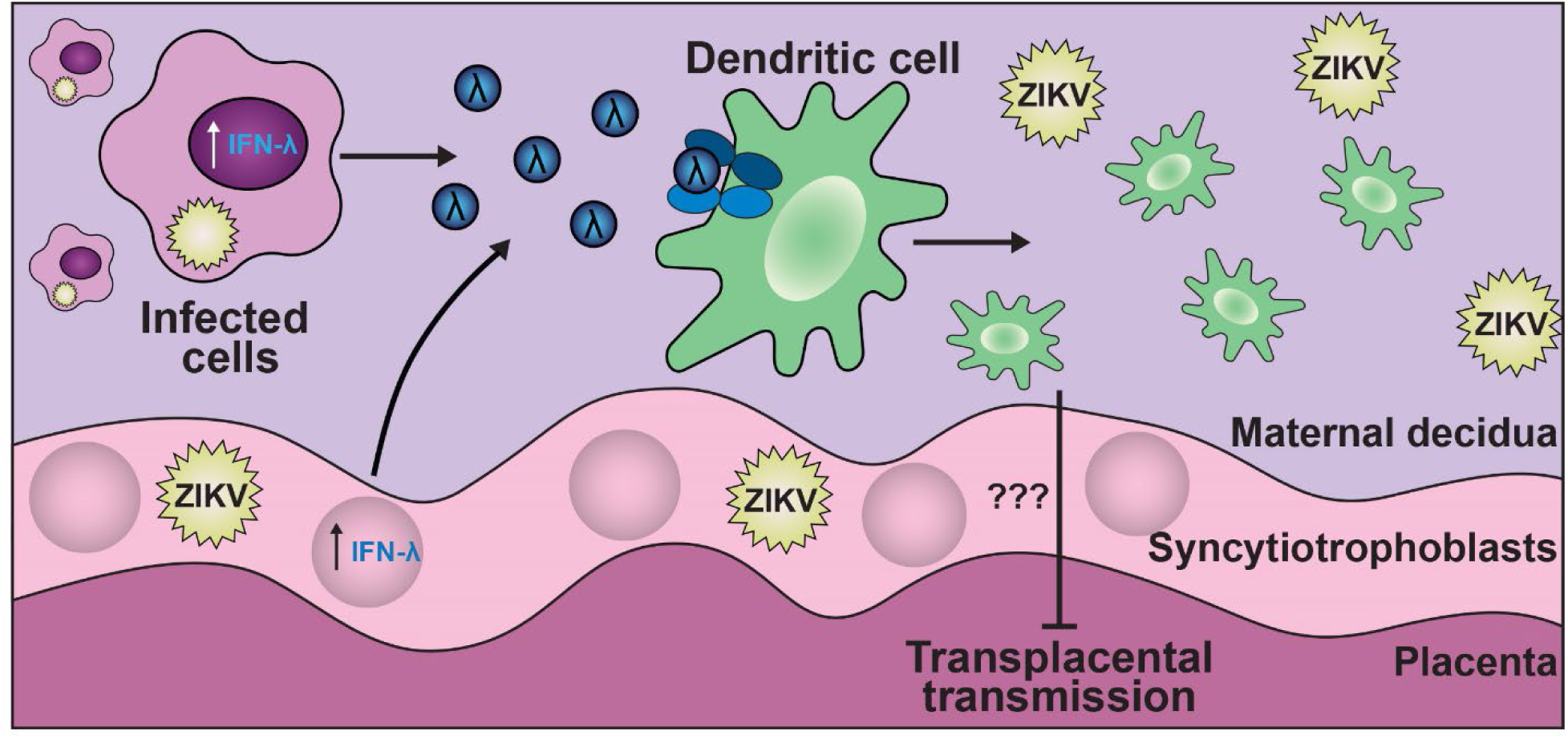
Proposed mechanism by which maternal IFN-λ signaling protects against transplacental transmission during congenital ZIKV infection. Infected cells produce IFN-λ, which signals to maternal dendritic cells. Dendritic cells infiltrate the maternal-fetal interface and block fetal infection through an unknown mechanism.

*LysM*-Cre-*Ifnlr1^-/-^* dams exhibited significantly higher fetal viral loads compared to Cre-dams (median 3.37 vs. 1.94 Log_10_ copies/mL, *P* < 0.05) and similar fetal viral loads compared to *Actin*-Cre-*Ifnlr1^-/-^* dams (median 3.37 vs. 4.07 Log_10_ copies/mL). Dams lacking IFN-λ signaling in myeloid cells (*LysM*-Cre-*Ifnlr1^-/-^*) exhibited an increased rate of transplacental transmission (73% ZIKV positive) similar to that observed in mice lacking IFN-λ signaling globally, suggesting that IFN-λ signaling to maternal myeloid cells promotes lower fetal viral loads and decreased transplacental transmission. *Mrp8*-Cre-*Ifnlr1^-/-^* dams exhibited no significant difference in fetal viral loads compared to Cre-dams (median 2.91 vs. 1.94 Log_10_ copies/mL) or fetal infection rates (43% vs. 56%), suggesting that neutrophils do not mediate IFN-λ-dependent protection against fetal infection. In contrast, dams lacking IFN-λ signaling in dendritic cells *CD11c*-Cre-*Ifnlr1^-/-^* exhibited increased fetal viral loads (median 4.59 vs. 1.94 Log_10_ copies/mL, *P* < 0.0001) and fetal infection rates (89% vs. 56%, *P* < 0.0001). Fetal viral loads and infection rates in these mice were comparable to those observed in *Actin*-Cre-*Ifnlr1^-/-^* dams, suggesting that maternal dendritic cells are the key IFN-λ responsive cell type mediating protection against ZIKV congenital infection.

## DISCUSSION

IFN-λ has an important role in combating viral infections at anatomic barriers by inducing an antiviral response, maintaining barrier integrity, and modulating the response of immune cells, which altogether promotes a local response that is less inflammatory and damaging than that of type I interferons^33,36,38,39,48–53^. IFN-λ has a well-documented role at the maternal-fetal interface during congenital infections^20,41–44,54,55^, but the specific cell types that respond to IFN-λ to mediate its protective and pathogenic effects at this site are not defined. To investigate the role of maternal IFN-λ signaling in protection against congenital ZIKV infection, we used a variety of transgenic mouse lines that lack IFN-λ signaling. Since our prior work showed that IFN-λ acted specifically through signaling in maternal tissues^44^, here we set up pregnancies such that all fetuses and placentas retained IFN-λ signaling, allowing us to focus specifically on the effects of maternal IFN-λ signaling. We found that IFN-λ restricts ZIKV congenital infection, as dams that lack IFN-λ signaling (*Ifnar1^-/-^ Ifnlr1^-/-^*) had increased fetal viral loads and transplacental transmission rates, regardless of the duration of the infection. The protective effects of IFN-λ were specific to transplacental transmission because maternal IFN-λ signaling had no impact on infection in maternal tissues, consistent with prior findings demonstrating a lack of effect of IFN-λ on systemic infection^42,44,50,56^. Although RT-qPCR did not reveal IFN-λ-dependent changes in antiviral gene responses in placenta homogenates, microscopy analysis of whole gravid uteruses probed by HiPlex RNAscope revealed that IFNs were expressed more proximally to sites of ZIKV infection in dams with intact IFN-λ signaling, implying that maternal IFN-λ signaling facilitates a protective antiviral response in the placenta which may restrict viral infection to localized regions of the placenta, preventing transplacental transmission. These findings highlight the importance of spatial relationships between infection and immune responses in the complex architecture of the maternal-fetal interface. Using flow cytometry, we found that a lack of maternal IFN-λ signaling resulted in a decrease in NK cells and dendritic cells in the uterus. Comparison of fetal infection rates in *Ifnlr1^-/-^* conditional knockout mice revealed that maternal myeloid cells, specifically dendritic cells, are key IFN-λ responsive cell types required for protection against fetal infection. Altogether, our data support a model where maternal IFN-λ signaling to dendritic cells increases recruitment to the uterus and facilitates a protective antiviral response that restricts ZIKV fetal infection (Figure 7).

In dams that lack IFN-λ signaling, we observed increased fetal viral loads and transplacental transmission compared to dams with intact IFN-λ signaling, but these higher fetal viral loads did not correspond to increased fetal pathology. In this study, we used *Ifnar1*^-/-^ dams and sires to achieve more robust ZIKV infection than is sustained in mice treated with an IFNAR1-blocking mAb^57^, but this mating scheme results in *Ifnar1*^-/-^ fetuses and placentas. Since fetal and/or placental IFN-αβ signaling is a significant driver of fetal pathology and resorptions^44,45^, the relatively low levels of fetal pathology observed in our experiments are not unexpected. Furthermore, in our model, observable fetal pathology is limited to low fetal weights (IUGR) or resorptions, which does not account for placental pathology, more subtle fetal developmental defects, or long-term neurological defects. Congenital ZIKV in mice can result in impaired angiogenesis, neurological defects including brain and spinal cord malformations, and subsequent neurological disorders including paralysis and behavioral changes^58–63^, consistent with outcomes evident in children infected with ZIKV in utero^3,64–66^. It is possible that higher fetal viral loads correlate with one or more of these known outcomes of congenital ZIKV infection in mice, even in the absence of IUGR or resorption.

Since ZIKV is a mosquito-borne virus that causes viremia in humans and animal models, a straightforward mechanism for the virus to spread to the fetal compartment would be hematogenous transplacental transmission, spreading from maternal blood, across the placental interface, into the fetal circulation. However, demonstrating the actual route of fetal infection in humans or animal models is challenging^67^. In pregnant mouse models, dissected placentas typically retain closely associated decidual tissue, so detection of ZIKV RNA in placental homogenates cannot distinguish true placental infection from infection of the maternal decidua; this is particularly important in experiments using *Ifnar1*^-/-^ dams, which sustain high levels of viral infection. ZIKV infection of the placenta has been detected by immunohistochemistry and RNA hybridization approaches in mice and macaques^46,47,61,63^, but this could occur secondary to fetal infection by other routes. Histologic analysis of infected mouse placentas often removes this tissue from the uterus prior to processing, precluding an understanding of how placental infection corresponds to infection of other tissues at the maternal-fetal interface. In this study, we embedded entire gravid uteruses for histologic analysis, preserving the spatial architecture of the fetal compartment within the uterus. This method required pregnancies to be harvested at E13, because at our standard harvest time of E15, the gravid uterus is too large to fit into a standard tissue cassette, and the more extensive fetal development produced poorer-quality tissue sections. Using RNAscope, we detected ZIKV RNA in the uterus, placenta, and fetus, but the most abundant ZIKV RNA signal was detected in the uterus, identifying this tissue as a key site of ZIKV infection at the maternal-fetal interface, and consistent with RT-qPCR data showing that the uterus has higher viral loads per gram compared to the placenta or fetus. Notably, a study in a rhesus macaque model of congenital ZIKV infection detected robust infection in the uterus and fetal membranes, rather than the placenta, supporting a model wherein ZIKV spreads to the fetal compartment by a route that bypasses the placenta^46,47^. Such a route would be consistent with findings that the syncytiotrophoblasts that form the interface with maternal blood are highly resistant to infection, including by ZIKV, in part due to constitutive expression of IFN-λ^40,41,68^. Our detection of abundant ZIKV RNA signal in the uterus supports the idea that there are additional routes of fetal infection beyond hematogenous transplacental transmission.

We found that *Ifnar1^-/-^ Ifnlr1^-/-^* dams have decreased NK cell and dendritic cell accumulation in the uterus compared to *Ifnar1^-/-^* dams, suggesting that maternal IFN-λ signaling promotes accumulation or infiltration of these cells at the maternal-fetal interface during congenital ZIKV infection. This effect was not perfectly recapitulated in the placenta, where *Ifnar1^-/-^ Ifnlr1^-/-^*dams exhibited a decrease in NK cells and dendritic cells as a percentage of leukocytes, but not in the total number of cells. Interpretation of flow cytometry results from dissociated placentas is somewhat confounded by decidual tissue that remains closely associated with the placenta when this tissue is dissected from the gravid uterus, meaning that placental leukocyte preparations likely include decidual leukocytes as well. Since the decidua contains an abundance of leukocytes, particularly decidual NK cells^69–71^, the effect of maternal IFN-λ signaling on uterine NK cells and dendritic cells may contribute to differences observed in the placentas of these mice. Future studies will incorporate congenic markers to more readily distinguish maternal and fetal leukocytes in these mixed-source tissues.

Although we found that maternal IFN-λ signaling corresponded to higher NK cell accumulation in the uterus, when we depleted maternal NK cells we found no effect on fetal infection, arguing against a key role for NK cells in IFN-λ-mediated protection against ZIKV congenital infection. For this experiment, we administered ⍺NK1.1 antibody at the time of infection (E9), and every two days until harvest (E15), resulting in a decrease in NK cells in the blood, spleen, uterus, and placentas as assessed by flow cytometry. Notably, while leukocyte depletions by administration of antibodies have been performed in pregnant mice^72–76^, and have been shown to deplete target cells in the pregnant uterus^77^, to our knowledge, it had not previously been demonstrated that this approach successfully depletes target cells in the placenta; our results demonstrate that this approach can be used to further investigate immune-mediated protection in the placenta, as well as in maternal tissues. While we observed successful depletion of NK cells at the maternal-fetal interface when tissues were harvested at E15, it remains possible that at the time of infection, or throughout infection, there were sufficient NK cells present at the maternal-fetal interface to contribute to protection against fetal infection. Additionally, while NK cells may not have an important role in protection against fetal infection in this model, it is possible that NK cells accumulate at the maternal-fetal interface in a maternal IFN-λ signaling dependent manner for other purposes. NK cells are critical cell types during pregnancy and play important roles in development of the fetus and placenta including aiding in the formation of the placenta and spiral artery, contributing to immune tolerance, and promoting fetal development^78,79^. NK cells may infiltrate the maternal-fetal interface during congenital ZIKV infection to promote immune tolerance in the context of the robust anti-viral response promoted by maternal IFN-λ signaling.

Our results show that dendritic cells accumulate at the maternal-fetal interface and protect against ZIKV fetal infection in a manner dependent on maternal IFN-λ signaling, but the mechanism by which dendritic cells mediate protection against fetal infection remains unknown. The IFN-λ dependent accumulation of dendritic cells at the maternal-fetal interface may promote a more robust antiviral response, perhaps one more optimally localized to specific sites of ZIKV infection. Circulating myeloid cells, including dendritic cell are key targets for ZIKV infection, with these cells often serving as some of the first cells to become infected in the skin and the genital tract^80–82^. It is possible that dendritic cells at the maternal-fetal interface are infected with ZIKV, but maternal IFN-λ signaling to these cells produces an antiviral response that limits spread of the virus beyond infected cells. Furthermore, IFN-λ is known to alter dendritic cell function and the interaction between dendritic cells and CD8 T cells in the context of influenza A virus and SARS-CoV-2 infections^83,84^. Although we did not observe differences in CD4 or CD8 T cell accumulation in the absence of maternal IFN-λ signaling, it is possible that IFN-λ signaling to dendritic cells could impact the quality of antiviral CD8 T cell responses at the maternal-fetal interface.

Our group and others have demonstrated that neutrophils are a key IFN-λ responsive cell type in mice in a variety of infection and inflammatory contexts^28,53,85–87^. However, our experiments using conditional knockout mice found that dams lacking IFN-λ signaling in neutrophils (*Mrp8*-Cre-*Ifnlr1*^-/-^) had similar fetal viral loads and transplacental transmission rates as dams with intact IFN-λ signaling, indicating that neutrophils are not key IFN-λ responsive cells mediating protection of the fetus during congenital ZIKV infection. The effect of maternal IFN-λ signaling on neutrophil accumulation in the uterus and placenta was difficult to assess in our flow cytometry experiments, due to technical obstacles to distinguishing neutrophils in the circulation from neutrophils in tissue; future studies will incorporate *in vivo* intravascular staining approaches to better assess neutrophil abundance and activity at the maternal-fetal interface.

Altogether, our data provide the first evidence for regulation of dendritic cell function by IFN-λ signaling at the maternal-fetal interface during congenital infection. Using a combination of spatial analysis and immune phenotyping approaches, we have shown that maternal IFN-λ signaling limits ZIKV transplacental transmission, amplifies IFN-λ expression at the site of infection, and promotes the accumulation of NK cells and dendritic cells at the maternal-fetal interface during congenital ZIKV infection. Our results further highlight the importance of the maternal immune response during congenital infections.

## MATERIALS AND METHODS

### Cells and viruses

Vero (African green monkey kidney epithelial) cells were maintained in Dulbecco’s modified Eagle medium (DMEM) containing 5% heat-inactivated fetal bovine serum (FBS) and 1X GlutaMAX (Gibco # 35050061) at 37°C with 5% CO_2_. Virus stocks were grown in Vero cells in DMEM containing 2% FBS and 10mM HEPES at 37°C with 5% CO_2_. ZIKV strain FSS13025 (Cambodia 2010) was obtained from the World Reference Center for Emerging Viruses and Arboviruses^88^. Virus stock titers were determined by focus forming assay using anti-flavivirus human monoclonal antibody E60^89^ at 100ng/ml (purified from hybridoma supernatant by the UNC Protein Expression Core Facility), HRP-conjugated goat anti-mouse antibody at a 1:5000 dilution (Fisher # 50-673-91), and TrueBlue Peroxidase substrate (KPL). Antibody incubations were performed for 2 hours at room temperature. Foci were counted on a CTL Immunospot analyzer.

### Mice

All experiments and husbandry protocols were approved by the Institutional Animal Care and Use Committee at the University of North Carolina at Chapel Hill. Experiments used 8-to 20-week-old female mice on a C57BL/6 background mated with 8-to 52-week-old male mice on a C57BL/6 background. *Ifnar1^-/-^* mice and *Ifnar1^-/-^ Ifnlr1^-/-^* mice were bred in-house as knockout x knockout^44^. *Actin-*Cre-*Ifnlr1^-/-^*, *LysM-*Cre-*Ifnlr1^-/-^*, *Mrp8-*Cre-*Ifnlr1^-/-^*, and *CD11c-*Cre-*Ifnlr1^-/-^* mice were generated by crossing *Ifnlr1^f/f^* mice with mice expressing Cre recombinase under control of the *Actin* promoter (Jackson labs #19099; obtained from Jenny Ting, UNC Chapel Hill), *LysM* promoter (Jackson labs #4781; obtained from Jenny Ting, UNC Chapel Hill), *Mrp8* promoter (Jackson labs #21614), or *Cd11c* promoter (Jackson labs #8068; obtained from Jenny Ting, UNC Chapel Hill) ^53^. Mice were bred as Cre hemizygotes, generating mixed litters in which 50% of mice were Cre+, lacking IFN-λ signaling in respective cell types, and 50% of mice were Cre-, retaining IFN-λ signaling. Experiments were performed blinded to Cre status, with tail snips from experimental mice being genotyped after data collection by PCR for both Cre and *Ifnlr1*.

### Genotyping

For conditional knockout mice, *Ifnlr* and Cre genotypes were determined by PCR on tail samples following harvest of pregnant dams. DNA was extracted from tails using Quantbio DNA extraction kit (#95091-250). PCR was run using Accustart II Geltrack PCR Supermix (#89235-014) and the following primers:

*Ifnlr*: F^1^ 5’-AGGGAAGCCAAGGGGATGGC-3’, R^1^ 5’-AGTGCCTGCTGAGGACCAGGA-3’, R^3^ 5’-GGCTCTGGACCTACGCGCTG-3’

*Actin*-Cre: F^1^ 5’-CGTACTGACGGTGGGAGAAT-3’, R^1^ 5’-CCCGGCAAAACAGGTAGTTA-3’ *Cd11c*-Cre: F^1^ 5’-ACTTGGCAGCTGTCTCCAAG-3’, R^1^ 5’-GCGAACATCTTCAGGTTCTG-3’ *LysM*-Cre: F^1^ 5’-CTTGGGCTGCCAGAATTTCTC-3’, R^1^ 5’-CCCAGAAATGCCAGATTACG-3’, R^2^ 5’-TTACAGTCGGCCAGGCTGAC-3’

*Mrp8*-Cre: F^1^ 5’-GCGGTCTGGCAGTAAAAACTATC-3’, R^1^ 5’-GTGAAACAGCATTGCTGTCACTT-3’

For *Actin*-Cre, *Cd11c*-Cre, and *Mrp8*-Cre, genotyping, two primers (F^1^ 5’-CAAATGTTGCTTGTCTGGTG-3’, R^1^ 5’-GTCAGTCGAGTGCACAGTTT-3’) were used to amplify an internal positive control.

### Mouse experiments

Females were exposed to soiled male bedding 3 days prior to mating to promote estrus. Male and female pairs were co-housed overnight (E0) and were separated in the morning (E1). Female mice were inoculated via subcutaneous injection in the footpad with 1000 FFU of ZIKV in 50µl 9 days post-mating (E9). Conditional knockout mice were administered 2mg of anti-IFNAR1-blocking antibody MAR1-5A3 via intraperitoneal injection 1 day prior to infection (E8).^57^ Pregnant mice were harvested 6 days post-infection (E15) or 8 days post-infection (E17). Cardiac puncture was used to harvest maternal blood in serum separator tubes (BD). Serum was separated from whole blood by centrifugation at 8,000 rpm for 5 minutes. Fetuses, placentas, and maternal spleens were harvested and weighed. Photographs of fetuses and placentas were taken at time of harvest. Any implantation without discrete fetal and placental structures was considered a resorption. For microscopy experiments, pregnant mice were harvested 4 days post-infection (E13). Dams were perfused with 20 mL of PBS and then the entire gravid uterus was harvested, or fetuses, placentas, and maternal uteruses were harvested and weighed.

### Viral genome quantification by RT-qPCR

Tissues were homogenized in 1mL PBS using a MagNA Lyser (Roche) and equal parts homogenate was added to equal parts RLT buffer (Qiagen) (150µL for fetuses and placentas, 300µL for spleens and uteruses). Viral RNA was extracted using a Qiagen RNeasy kit and detected by TaqMan one-step RT-qPCR using the following primer/probe set: forward-TTGGTCATGATACTGCTGATTGC; reverse-CCTTCCACAAAGTCCCTATTGC; probe56-FAM/CGGCATACA/ZEN/GCATCAGGTGCATAGGAG/3IABkFQ on a Bio-Rad CFX96. ZIKV copies per mL were determined by comparison to a standard curve of 10-fold dilutions of ZIKV-A plasmid.^90^

### Plaque assay

Tissues were homogenized in 1mL PBS and silicone beads using a MagNA Lyser (Roche) and serially diluted in a round bottom 96-well plate. Beginning with the undiluted sample, 100µL of each dilution and was added to monolayers of Vero cells (6-well plate with 5 x 10^5^ cells/well plated 1 day prior) with 500µL of DMEM with 2% FBS and 10mM HEPES, and were incubated for 1 hour at 37°C. After 1 hour, media was removed and 2mL methylcellulose (1% methylcellulose in 1X MEM + 2% FBS, L-Glut, P/S, HEPES) was added to the wells and left to incubate at 37°C for 5 days. After 5 days, media was aspirated, and the plates were fixed by adding 500µL of 10% formaldehyde for at least 30 minutes at room temperature. After fixing, PFA was poured off, plates were washed with tap water, stained with crystal violet (1% crystal violet, 20% ethanol, 80% water), and washed again with tap water. Plaques were counted after plates dried at room temperature overnight.

### HiPlex RNAscope

Entire gravid uteruses were harvested and fixed in 10% neutral buffered formalin for 72 hours. Tissues were processed using a Sakura Tissue-Tek VIP 5 Tissue Processor, embedded in paraffin, and sectioned into 5µm slices. *In situ* hybridization was performed using ACD Bio’s RNAScope technology per the kit instructions (RNAScope Kit HiPlex 12 Detection kit V2 [488, 550, 650, 750] #324400). Briefly, tissue sections were deparaffinized and hydrated, followed by antigen retrieval for 15 minutes at 100°C. Tissue sections were incubated with protease for 30 minutes at 40°C. Then, probes were hybridized for two hours at 40°C. Following hybridization, probe signal was amplified. The following probes were used: V-ZIKV-T1 (#467771), Mm-Ifnl2-T2 (#410231), Mm-Ifnb1-T3 (#406531), Mm-Ifng-T4 (#311391), Mm-Ifit1-T5 (#500071), Mm-ITGAM-T6 (#311491), Mm-ITGAX-T7 (#311501), Mm-Cxcl9-T8 (#489341), Mm-Tpbg-T9 (#425681), Mm-Ifnlr1-T10 (#512981), Mm-Cd244-T11 (#486801), and Mm-Ngp-T12 (#1041701).

Fluorophores for T1-T4 were developed in the first round, then cover slips were added to slides using Fluorogel II with DAPI (Electron Microscopy Sciences, #17985-50). Tissue sections were imaged, then according to kit instructions, coverslips were removed, fluorophores for T1-T4 were cleaved, fluorophores for T5-T8 were developed, and coverslips were added. Tissue sections were imaged and the coverslip removal, fluorophore cleavage, and subsequent development process was repeated for T9-T12. Following imaging of T9-T12, fluorophores were cleaved, coverslips were added to slides, and sections were imaged without any fluorophores as a control for background fluorescence.

### Microscopy and image analysis

The samples were imaged following each round of probe hybridization using a using a Nikon Eclipse Ti2-D-LHLED (Nikon #MEE55700) with a linear encoded mechanical stage (Nikon #MEC56130). All images were taken with an air immersion 4x Plan Apo objective with a numerical aperture of 0.2 (Nikon #MRD00045). The following filters and exposure times were used for detection of fluorescent signal: GFP (Chroma 49002-ET-GFP, exposure time 400ms), TexasRed (Chroma 49008-ET-mCherry, Texas Red, exposure time 600ms), Narrow Cy5 (Chroma 49009-ET-Cy5 Narrow Excitation, exposure time 400ms), DAPI (Chroma NC764537-49000-ET-DAPI, exposure time 400ms), and Cy7 (Chroma 96367-ET-Cy7, exposure time 400ms). All filter cubes were hard coated. Filter cubes were in a motorized filter cube turret (Nikon Ti2-F-FLT-E, #MEV51030). Images were detected with a pco.edge 4.2 Q High sCMOS camera (Pco #77067008). This camera has a 16-bit maximum bit depth. NIS Elements is the acquisition software. Images were collected across the entire tissue section on the slide, stitched in NIS Elements (version AR), denoised using Nikon Batch Denoiser and subsequently converted to Imaris files. Images of each round of hybridization on the same sample were registered using the SITK-IBEX based registration package in Python^91^. Using Imaris (version 10.2.0) spot-detector, spots were assigned to the signal from probes based on the quality of the signal with the following parameters: ZIKV, 170.0; *Ifnl2/3,* 163.0; *Itgax* 25.0; and *CD244*, 21.5. Shortest distance between spots of interest was calculated in Imaris. Images shown are representative.

### Cellular gene expression measured by RT-qPCR

Gene expression was measured by extracting RNA from placental tissue as previously described, followed by cDNA synthesis with the Verso cDNA Synthesis Kit (Thermo Scientific). Reactions were set up according to the manufacturer’s instructions. Quantification of gene expression was performed using Power SYBR Green PCR master mix (Applied Biosystems) using the following primer combinations on a Bio-Rad CFX96: *Actb* (Fwd-CCTCTATGCCAACACAGTGC; Rev-CCTGCTTGCTGATCCACATC), *Ifnl3* (Fwd-GACAAGAACCCAAGCTGACC; Rev-ATGTCCTTCTCAAGCAGCCT), *Ifnb1* (Fwd-GTCCTCAACTGCTCTCCACT; Rev-CCTGCAACCACCACTCATTC), *Ifit1* (Fwd-GATGGACTGTGAGGAAGGCT; Rev-TGGATTTAACCGGACAGCCT).

### Flow cytometry

Tissues were harvested into cold PBS and processed using established methods for digestion of mouse uteruses^92^ and a modified method for digestion of mouse placentas. In brief, gravid uteruses were harvested at E15 and dissected to separate the uterus, fetuses, and placentas. Uteruses were cut into ∼0.5cm pieces and excess fat was trimmed and discarded. Then, uteruses were incubated in a predigestion solution (Hanks Balanced Salt Solution with 5% FBS, 5mM EDTA, and 1mM DTT) for two 30-minute periods at 37°C.

Intraepithelial leukocytes were collected after each period by passing uteruses through 70µm cell strainers. Intraepithelial leukocytes and uterus samples were combined and then dissociated using a GentleMACS (Miltenyi #130-134-029) and enzymes from the mouse lamina propria dissociation kit (Miltenyi #130-097-410). Uterus samples were passed through 70µm and 40µm filters, centrifuged at 350 x *g* for 5 minutes and resuspended in PBS with 1% FBS for staining with fluorescent antibodies. Separately, placentas were dissected into 8 pieces and washed with PBS. Placentas were then dissociated using a GentleMACS and enzymes from the mouse lung dissociation kit (Miltenyi #130-095-927). Samples were incubated at 37°C for 30 minutes and then dissociated again using the GentleMACS. Red blood cells were lysed using RBC lysis buffer (155mM NH_4_Cl, 12mM NaHCO_3_, 0.1mM EDTA), washed with PBS, and centrifuged at 500 x *g* for 5 minutes. Placental samples were passed through a 70µm cell strainer, centrifuged at 300 x *g* for 10 minutes and resuspended in PBS with 1% FBS (staining buffer), counted, and stained with fluorescent antibodies.

Spleens and whole blood were collected to confirm depletion of NK cells. Spleens were harvested into cold DMEM (Gibco) containing DNase I (100ug/mL, Fisher #NC9185812) and Collagenase P (1mg/mL, Sigma #11249002001) and then passed through 70µm filters and suspended in DMEM + 5% FBS. Red blood cells were lysed using ACK lysis buffer (Gibco #A1049201) and samples were washed with PBS, centrifuged at 300 x *g* for 5 minutes, and resuspended in staining buffer and stained with fluorescent antibodies. Whole blood was collected via cardiac puncture into tubes containing EDTA (Fisher #02-669-33). Red blood cells were lysed with ACK lysis buffer, centrifuged at 500 x *g* for 5 minutes, and resuspended in staining buffer for counting and staining with fluorescent antibodies.

Fc Block and all fluorescent antibodies used were sourced from BioLegend and titrated prior to experimental use. All samples were treated with Fc Block on ice for 10 minutes. Samples were washed with staining buffer and stained for surface antigens on ice for 30 minutes, washed three times with staining buffer, and fixed with 2% paraformaldehyde. Samples were analyzed on an Attune Nxt (ThermoFisher) and data analyzed with FlowJo Software (v10.10). The following antibodies were used: ⍺CD3 (17A2), ⍺CD4 (GK1.5), ⍺CD8 (53-6.7), ⍺CD11b (M1/70), ⍺CD11c (N418), ⍺CD19 (6D5), ⍺CD45 (30-F11), ⍺F4/80 (BM8), ⍺Ly6C (HK1.4), ⍺Ly6G (1A8), ⍺MHCII (M5/114.15.2), ⍺NK1.1 (PK136), ⍺TCRγδ (GL3), and Zombie Aqua Fixable Dye. Gating strategies for NK cell and dendritic cells are shown in Figure S6. All gating strategies were based on published immunophenotyping methods.^93–95^ Fluorescence minus one controls and single-color controls were used for gating and compensation.

### NK cell depletion

To deplete NK cells, mice were injected intraperitoneally on E9, E11, and E13 with 100µg of ⍺NK1.1 antibody (Clone PK136, BioXCell). Depletion of neutrophils in blood, spleen, uterus, and placentas was confirmed by flow cytometry at the end of the experiment in each cohort of mice.

### Statistics

All statistics were calculated using Graphpad Prism 10 or R. Viral loads were analyzed using a Mann-Whitney *U* test. Transplacental transmission and fetal pathology rates were analyzed using a Cochran-Armitage test. Flow cytometry data was analyzed using an unpaired *t*-test.

## ACKNOWLEDGEMENTS

This work was supported by R01 AI139512 and R01 AI175708 (H.M.L.). H.M.L. holds an Investigators in the Pathogenesis of Infectious Disease Award from the Burroughs Wellcome Fund. M.R.D. and R.L.C. were supported by T32 AI007419. M.R.D. and R.L.C. were supported by Dissertation Completion Fellowships from the UNC Graduate School. T.R.R. was supported by the Michael P. and Jean W. Carter Research Fund administered by Honors Carolina and by a Summer Undergraduate Research Fellowship from the UNC Office for Undergraduate Research.

C.R.R. was supported by T32 GM152779 and a Gilliam Fellowship from HHMI. This work was supported by a Core Facilities Pilot Project Award from the UNC School of Medicine. Microscopy was performed at the UNC Neuroscience Microscopy Core (RRID:SCR_019060), supported in part by funding from the NIH-NICHD Intellectual and Developmental Disabilities Research Center Support Grant P50 HD103573. HiPlex RNAscope was performed at the UNC Histology Research Core. Flow cytometry was performed in the UNC Flow Cytometry Core Facility (RRID:SCR_019170), supported in part by P30 CA016086 Cancer Center Core Support Grant to the UNC Lineberger Comprehensive Cancer Center.

## Supplemental figure legends

**Figure S1.**
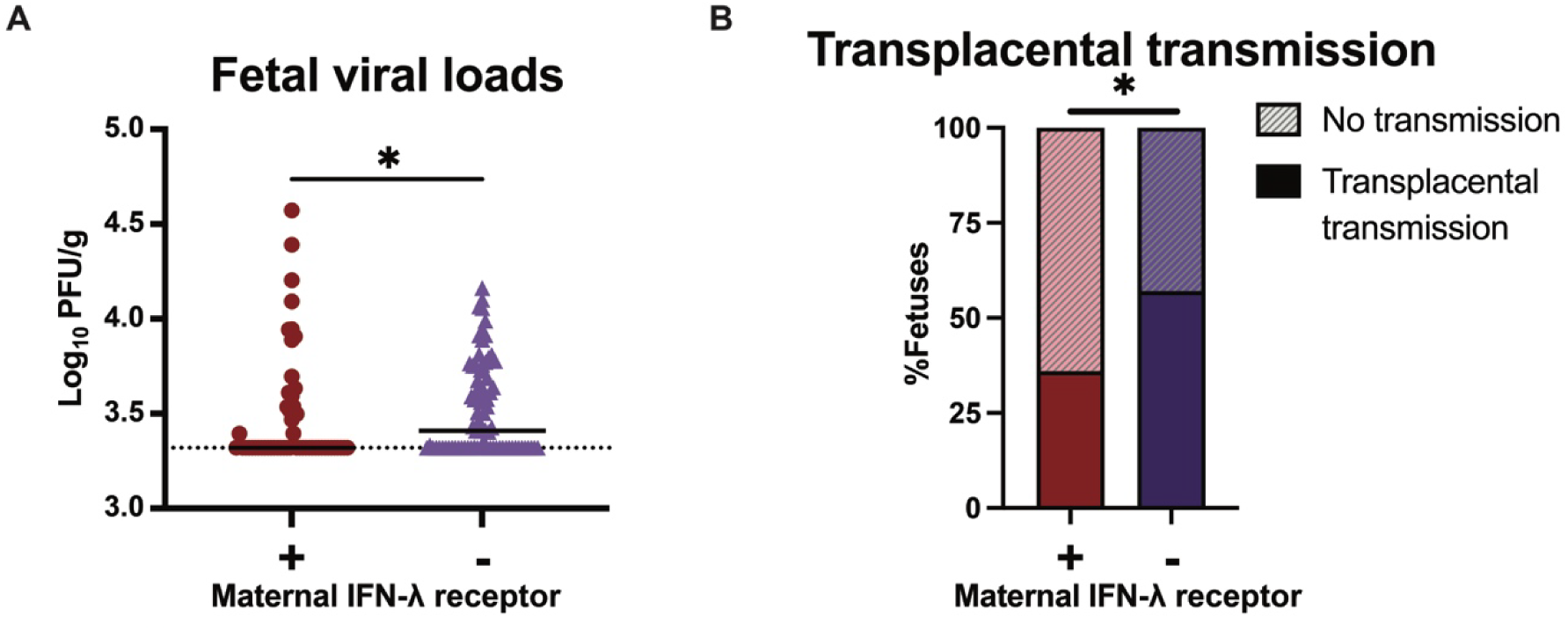
Maternal IFN-λ signaling results in decreased infectious viral titers in fetuses. Dams from *Ifnar1^-/-^* x *Ifnar1^-/-^ Ifnlr1^-/-^* or *Ifnar1^-/-^ Ifnlr1^-/-^* x *Ifnar1^-/-^* crosses were infected at day 9 post-mating (E9) with ZIKV FSS13025 by subcutaneous injection in the footpad. Fetal tissues were harvested at E15. (A) Viral loads in the fetus were measured by plaque assay. Each data point represents one fetus. (B) Transplacental transmission was calculated as percent of fetuses with viral loads above the limit of detection. Results are combined from 62 fetuses from 9 *Ifnar1*^-/-^ dams and 82 fetuses from 10 *Ifnar1^-/-^ Ifnlr1^-/-^* dams per group from 11 independent experiments. Viral loads were compared by Mann-Whitney test, proportions by Cochran Armitage test (*, *P* < 0.05).

**Figure S2.**
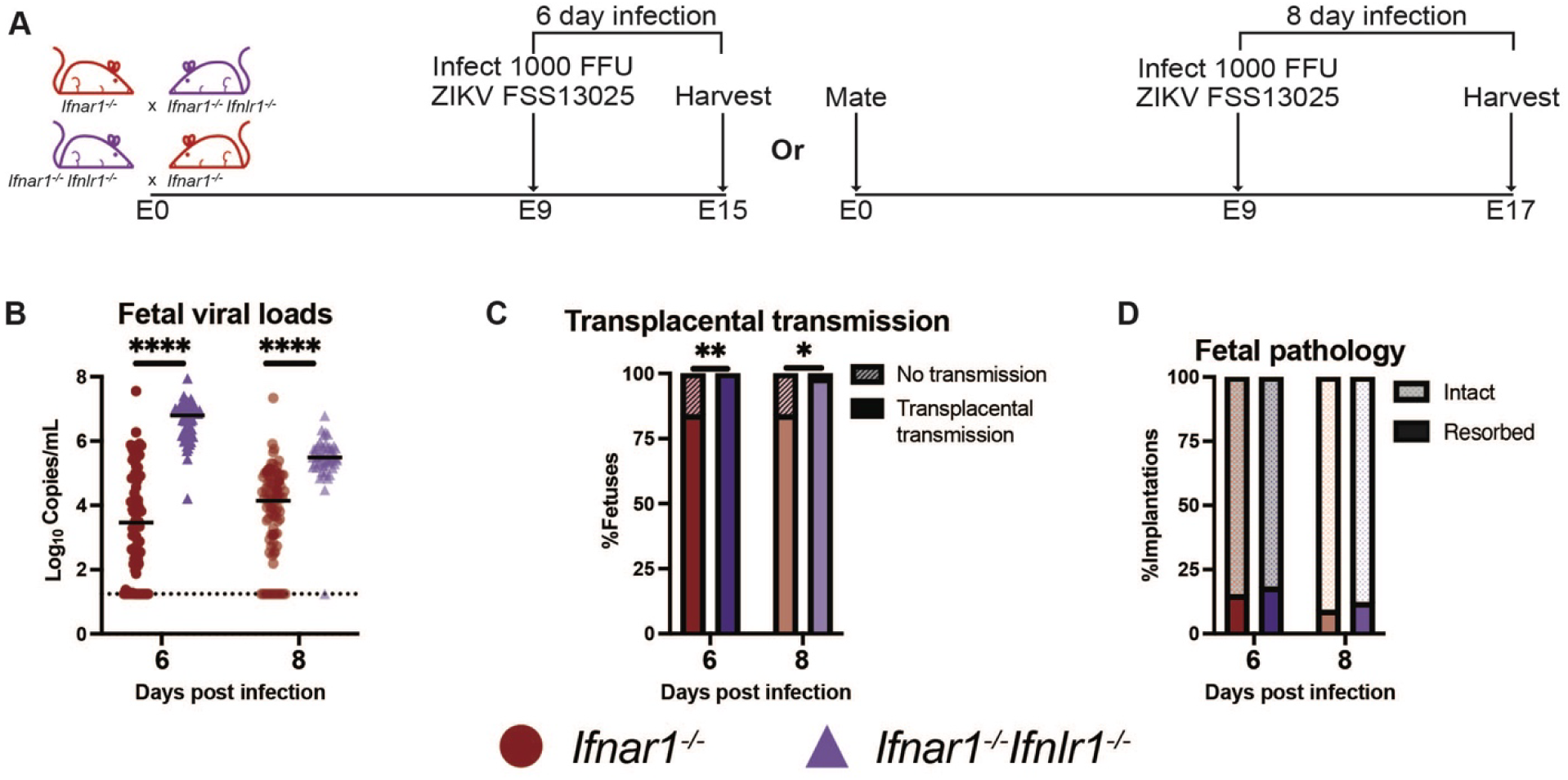
IFN-λ protects against fetal infection regardless of infection duration. (A) Dams from *Ifnar1^-/-^* x *Ifnar1^-/-^ Ifnlr1^-/-^* or *Ifnar1^-/-^ Ifnlr1^-/-^* x *Ifnar1^-/-^* crosses were infected at day 9 post-mating (E9) with ZIKV FSS13025 by subcutaneous injection in the footpad. Fetal tissues were harvested at E15 or E17. (B) ZIKV viral loads in the fetus were measured by RT-qPCR. Each data point represents one fetus. (C) Transplacental transmission was calculated as percent of fetuses with viral loads above the limit of detection. (D) Proportions of fetuses exhibiting resorption. Results are combined from 91 fetuses from 10 *Ifnar1*^-/-^ dams and 66 fetuses from 8 *Ifnar1^-/-^ Ifnlr1^-/-^* dams (E15 harvest) and 99 fetuses from 12 *Ifnar1*^-/-^ dams and 58 fetuses from 7 *Ifnar1^-/-^ Ifnlr1^-/-^* dams (E17). Viral loads were compared by Mann-Whitney test and proportions were compared by Cochran Armitage test (****, *P* < 0.0001; **, *P* < 0.01; *, *P* < 0.05).

**Figure S3.**
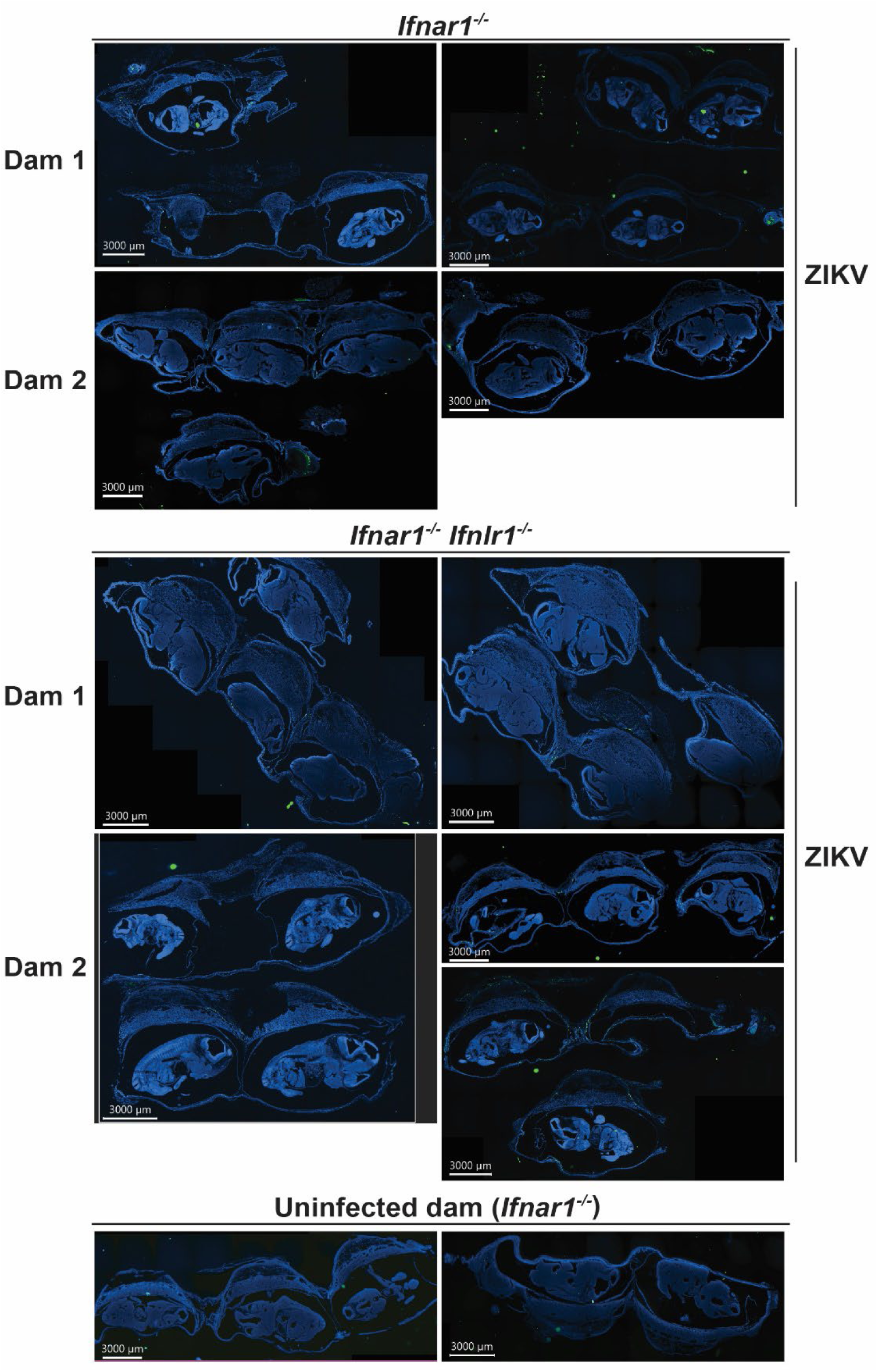
Sections of gravid uteruses. Dams from *Ifnar1^-/-^* x *Ifnar1^-/-^ Ifnlr1^-/-^* or *Ifnar1^-/-^ Ifnlr1^-/-^* x *Ifnar1^-/-^* crosses were infected at E9 with ZIKV FSS13025 by subcutaneous injection in the footpad. Gravid uteruses were harvested at E13. Sections of entire gravid uteruses (2 *Ifnar1*^-/-^dams and 2 *Ifnar1^-/-^ Ifnlr1^-/-^* infected dams, 1 uninfected *Ifnar1*^-/-^ dam, collected in independent experiments) were probed for ZIKV RNA (green) and stained with DAPI (blue).

**Figure S4.**
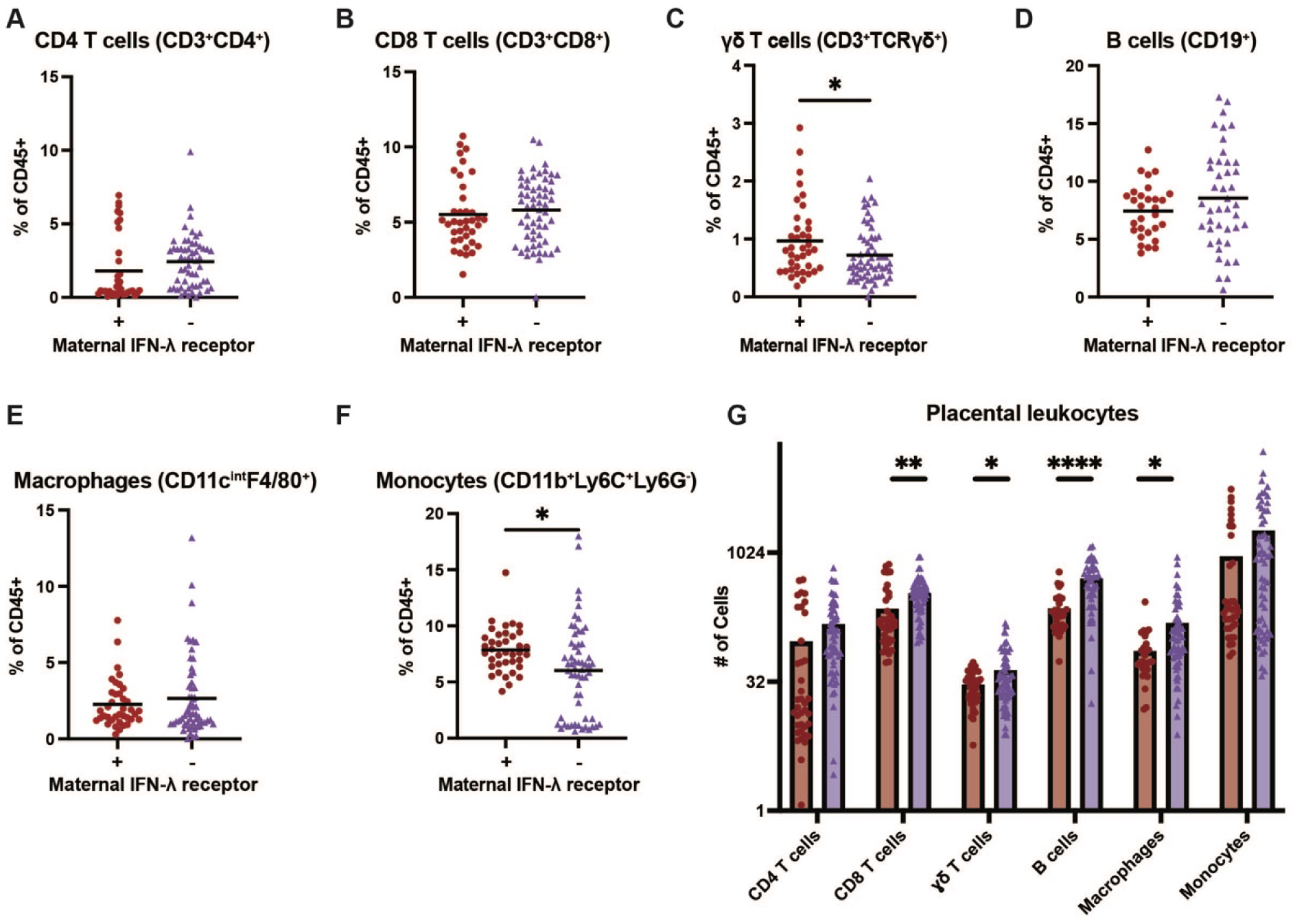
Immunophenotyping of placentas during ZIKV infection. Dams from *Ifnar1^-/-^* x *Ifnar1^-/-^Ifnlr1^-/-^* or *Ifnar1^-/-^ Ifnlr1^-/-^* x *Ifnar1^-/-^* crosses were infected at day 9 post-mating (E9) with ZIKV FSS13025 by subcutaneous injection in the footpad. Placentas were harvested at E15 and analyzed by flow cytometry (38 placentas from 5 *Ifnar1*^-/-^ dams and 59 placentas from 8 *Ifnar1^-/-^Ifnlr1^-/-^* dams collected in 6 independent experiments). (A) Frequency of CD4 T cells (CD3^+^CD4^+^ out of CD45^+^). (B) Frequency of CD8 T cells (CD3^+^CD8^+^ out of CD45^+^). (C) Frequency of ɣδ T cells (CD3^+^TCRɣδ^+^ out of CD45^+^). (D) Frequency of B cells (CD19^+^ out of CD45^+^). (E) Frequency of macrophages (CD11c^int^F4/80^+^ out of CD45^+^). (F) Frequency of monocytes (CD11b^+^Ly6C^+^Ly6G^-^out of CD45^+^). (G) Number of each leukocyte type quantified in placentas from *Ifnar1*^-/-^ and *Ifnar1^-/-^ Ifnlr1^-/-^* dams. Groups were compared by unpaired *t*-test (****, *P* < 0.0001; **, *P* < 0.01; *, *P* < 0.05).

**Figure S5.**
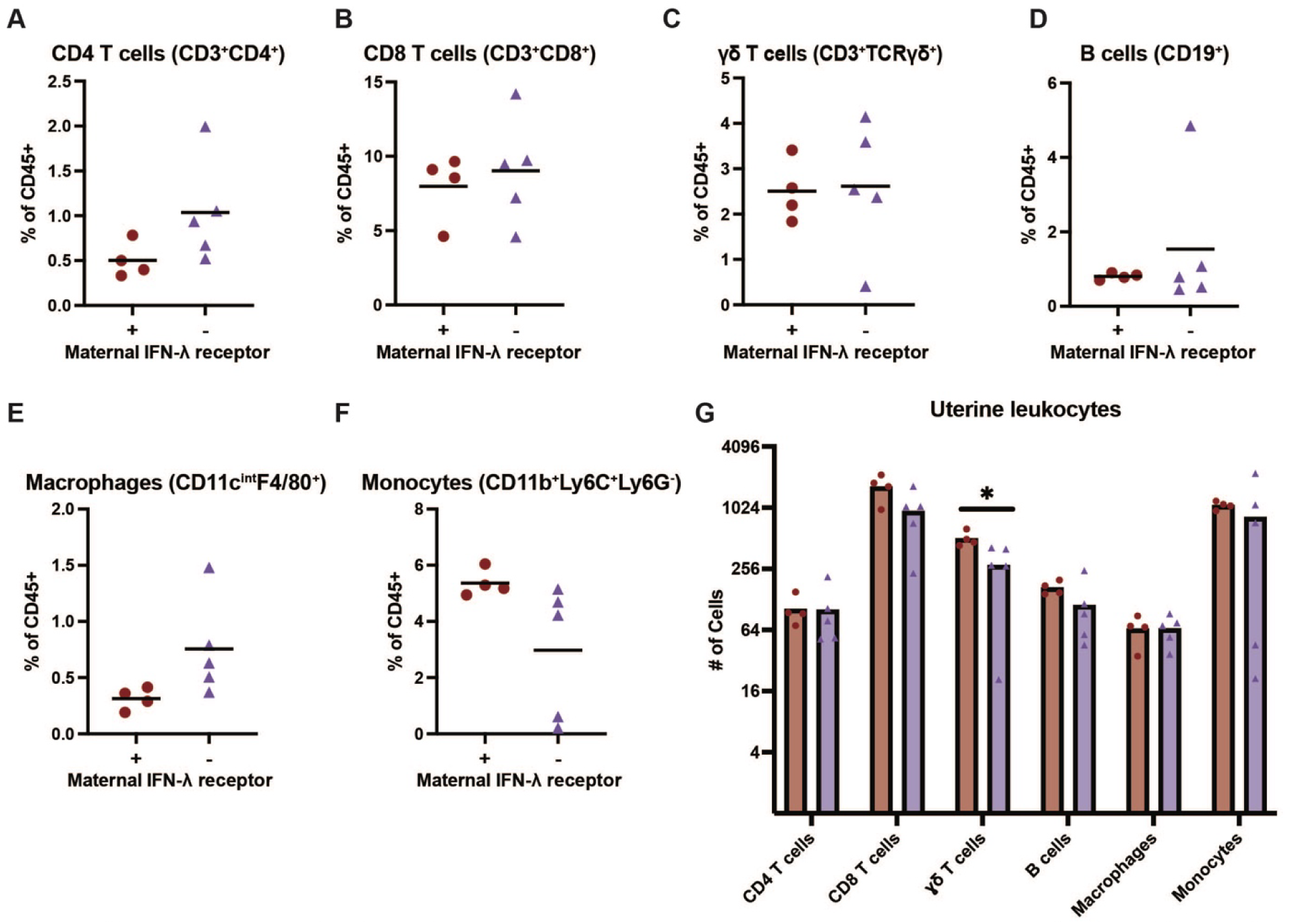
Immunophenotyping of the uterus during ZIKV infection. Dams from *Ifnar1^-/-^* x *Ifnar1^-/-^Ifnlr1^-/-^* or *Ifnar1^-/-^ Ifnlr1^-/-^* x *Ifnar1^-/-^* crosses were infected at day 9 post-mating (E9) with ZIKV FSS13025 by subcutaneous injection in the footpad. Uteruses were harvested at E15 and analyzed by flow cytometry (4 *Ifnar1*^-/-^ dams 5 *Ifnar1^-/-^ Ifnlr1^-/-^* dams collected in 4 independent experiments). (A) Frequency of CD4 T cells (CD3^+^CD4^+^ out of CD45^+^). (B) Frequency of CD8 T cells (CD3^+^CD8^+^ out of CD45^+^). (C) Frequency of ɣδ T cells (CD3^+^TCRɣδ^+^ out of CD45^+^). (D) Frequency of B cells (CD19^+^ out of CD45^+^). (E) Frequency of macrophages (CD11c^int^F4/80^+^ out of CD45^+^). (F) Frequency of monocytes (CD11b^+^Ly6C^+^Ly6G^-^ out of CD45^+^). (G) Number of each leukocyte type quantified in uteruses from *Ifnar1*^-/-^ and *Ifnar1^-/-^ Ifnlr1^-/-^* dams. Groups were compared by unpaired *t*-test (*, *P* < 0.05).

**Figure S6.**
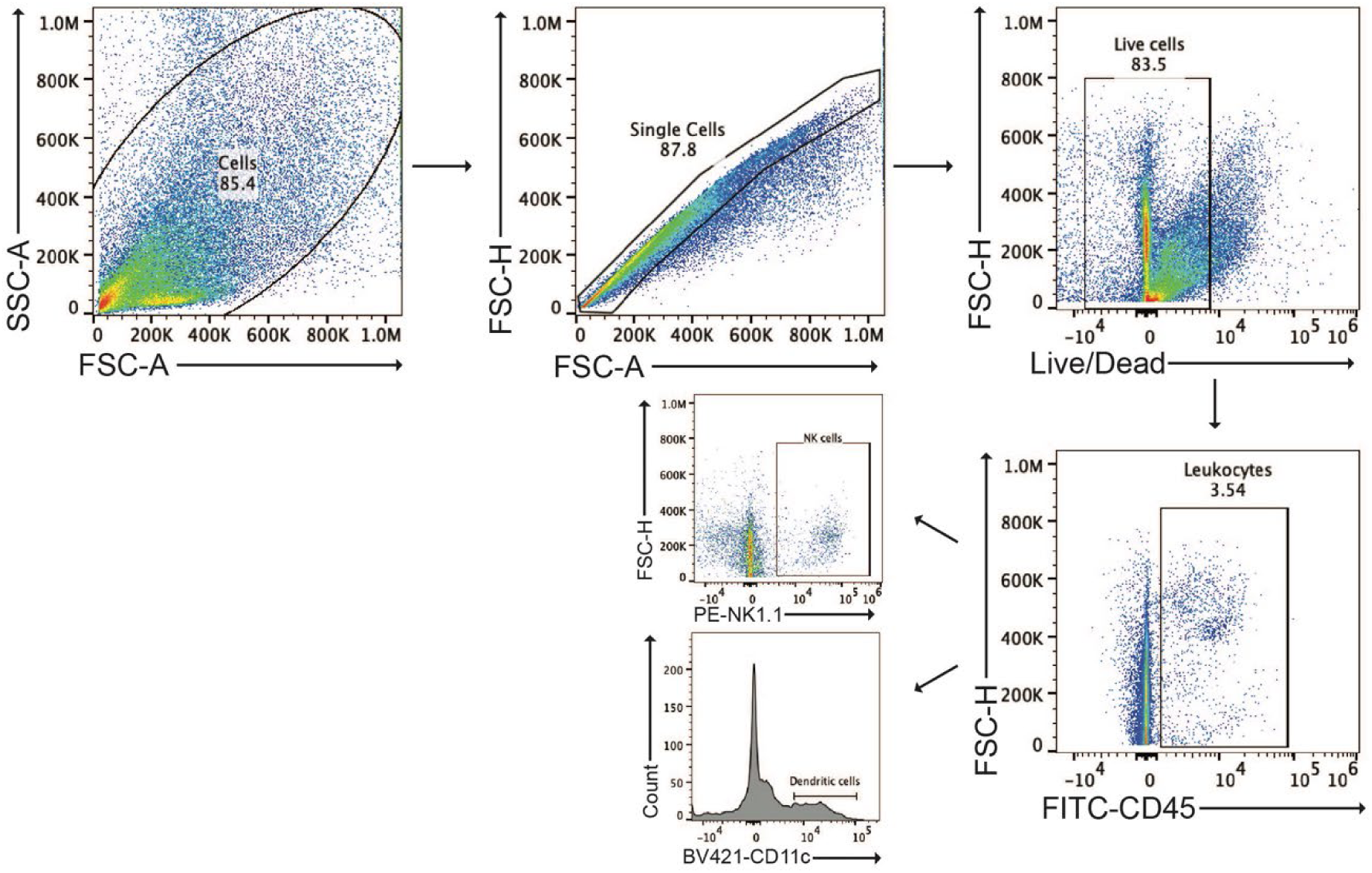
Representative gating strategy to define NK and dendritic cell populations. Gating strategy to define NK and dendritic cells in placentas/uteruses.

**Figure S7.**
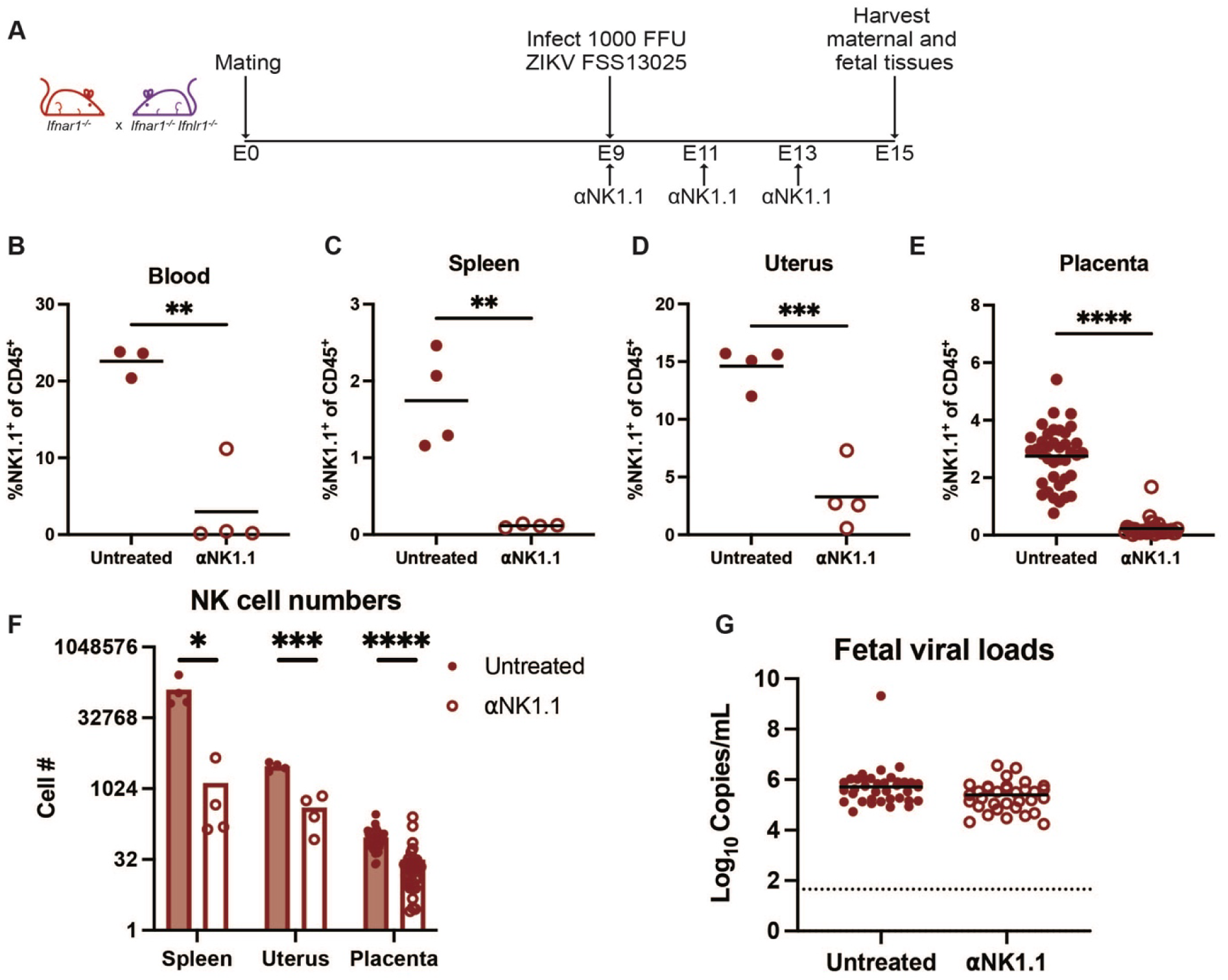
NK cells are not required for IFN-λ mediated protection against transplacental transmission. (A) Dams from *Ifnar1^-/-^* x *Ifnar1^-/-^ Ifnlr1^-/-^* crosses were infected at day 9 post-mating (E9) with ZIKV FSS13025 by subcutaneous injection in the footpad, and treated intraperitoneally at E9, E11, and E13 with 100µg of ⍺NK1.1 antibody. (B-F) Blood (3 untreated dams, 4 treated dams, collected in 4 independent experiments), spleens (4 untreated dams, 4 treated dams, collected in 4 independent experiments), uteruses ( 4 untreated dams, 4 treated dams, collected in 2 independent experiments), and placentas (38 placentas from 5 untreated dams, 33 placentas from 6 treated dams, collected in 6 independent experiments) were collected at E15 and analyzed for NK1.1^+^ cells by flow cytometry. Groups were compared by unpaired *t*-test (*, *P* < 0.05; **, *P* < 0.01; ***, *P* < 0.001; ****, *P* < 0.0001). (G) Fetuses (37 fetuses from 4 untreated dams, 33 fetuses from 4 treated dams, collected in 6 independent experiments) were collected at E15 for viral load analysis by RT-qPCR. Each data point represents one fetus. Groups were compared by Mann-Whitney and were found to have no significant difference in viral loads.

## Notes

### Competing Interest Statement

The authors have declared no competing interest.

